# Multi-parametric quantitative spinal cord MRI with unified signal readout and image denoising

**DOI:** 10.1101/859538

**Authors:** Francesco Grussu, Marco Battiston, Jelle Veraart, Torben Schneider, Julien Cohen-Adad, Timothy M. Shepherd, Daniel C. Alexander, Dmitry S. Novikov, Els Fieremans, Claudia A. M. Gandini Wheeler-Kingshott

## Abstract

Multi-parametric quantitative MRI (qMRI) of the spinal cord is a promising non-invasive tool to probe early microstructural damage in neurological disorders. It is usually performed by combining acquisitions with multiple signal readouts, which exhibit different thermal noise levels, geometrical distortions and susceptibility to physiological noise. This ultimately hinders joint multi-contrast modelling and makes the geometric correspondence of parametric maps challenging. We propose an approach to overcome these limitations, by implementing state-of-the-art microstructural MRI of the spinal cord with a unified signal readout. We base our acquisition on single-shot echo planar imaging with reduced field-of-view, and obtain data from two different vendors (vendor 1: Philips Achieva; vendor 2: Siemens Prisma). Importantly, the unified acquisition allows us to compare signal and noise across contrasts, thus enabling overall quality enhancement via Marchenko-Pastur (MP) Principal Component Analysis (PCA) denoising. MP-PCA is a recent method relying on redundant acquisitions, i.e. such that the number of measurements is much larger than the number of informative principal components. Here we used in vivo and synthetic data to test whether a unified readout enables more efficient denoising of less redundant acquisitions, since these can be denoised jointly with more redundant ones. We demonstrate that a unified readout provides robust multi-parametric maps, including diffusion and kurtosis tensors from diffusion MRI, myelin metrics from two-pool magnetisation transfer, and T1 and T2 from relaxometry. Moreover, we show that MP-PCA improves the quality of our multi-contrast acquisitions, since it reduces the coefficient of variation (i.e. variability) by up to 15% for mean kurtosis, 8% for bound pool fraction (BPF, myelin-sensitive), and 13% for T1, while enabling more efficient denoising of modalities limited in redundancy (e.g. relaxometry). In conclusion, multi-parametric spinal cord qMRI with unified readout is feasible and provides robust microstructural metrics with matched resolution and distortions, whose quality benefits from MP-PCA denoising, a useful pre-processing tool for spinal cord MRI.

## 1. Introduction

The spinal cord is a small but functionally important structure of the human central nervous system, affected in several common disorders. These are often associated with high disability (Hendrix et al., 2015), and include: multiple sclerosis (Ciccarelli et al., 2019), amyotrophic lateral sclerosis (van Es et al., 2017), spinal cord injury (Ahuja et al., 2017) and many others (Lorenzi et al., 2019). Routine anatomical magnetic resonance imaging (MRI) plays an important role in the diagnosis and management of these conditions (Kearney et al., 2015). However, it only offers macroscopic descriptors of tissue damage that lack specificity for pathophysiology, have limited prognostic value and fail to guide treatment and rehabilitation personalisation (Cohen-Adad, 2018; Stroman et al., 2014; Wheeler-Kingshott et al., 2014). The gradual adoption of quantitative MRI (qMRI) techniques may help overcome the limitations of conventional anatomical MRI. Based on either well-validated biophysical models or parsimonious signal representations (Novikov et al., 2018), qMRI promises to provide estimates of biologically meaningful characteristics, which would make parametric maps vendor-independent (Cercignani and Bouyagoub, 2018). The latest multimodal qMRI techniques exploit the complementary information from different contrasts (De Santis et al., 2016; Stikov et al., 2015), for example relaxometry and diffusion, to better quantify the parameters of tissue microstructure (Lemberskiy et al., 2018; Ning et al., 2019; Slator et al., 2019; Veraart et al., 2018).

qMRI of the spinal cord is increasingly popular (Battiston et al., 2018a; Battiston et al., 2018b; By et al., 2017, 2018; Duval et al., 2017; Grussu et al., 2019; Grussu et al., 2015; Ljungberg et al., 2017; Massire et al., 2016; Schilling et al., 2019; Taso et al., 2016) due to recent advancements in scanner hardware (Barry et al., 2018; Duval et al., 2015) and analysis software (De Leener et al., 2017). However, its development is currently hampered by the following two challenges.

Firstly, multi-contrast qMRI in the spinal cord typically relies on specialised techniques with dedicated signal readout for each contrast (Duval et al., 2017; Massire et al., 2016; Taso et al., 2016). The variety of readouts is not compatible with joint computational modelling of voxel-wise multi-contrast signals, and also limits the alignment of multimodal metrics due to different distortions and susceptibility to physiological noise (Campbell et al., 2018). The second major challenge is related to the fact that data quality in spinal cord imaging remains lower compared to the brain. This is due to the need for high spatial resolution (the spinal cord cross sectional area is about 1 cm^2^), which is challenged by artifacts from pulsation (Morozov et al., 2018; Summers et al., 2006) and local magnetic field inhomogeneities (Vannesjo et al., 2018; Verma and Cohen‐Adad, 2014). Improving the intrinsic quality of qMRI data is therefore imperative to facilitate the application of the latest qMRI techniques to the spinal cord, which are still in their infancy as compared to those in the brain (Cohen-Adad, 2018; Wheeler-Kingshott et al., 2014).

In this paper, we propose a unified acquisition for state-of-the-art multimodal qMRI of the spinal cord that addresses both challenges. Our protocol relies on a unified signal readout based on single-shot spin echo planar imaging (EPI) with reduced field-of-view (rFOV), which provides multi-contrast images, each having the same intrinsic resolution and susceptibility artifacts (i.e. distortions). Importantly, the unified acquisition enforces the same noise statistics across multiple signal contrasts, thus enabling overall data quality enhancement via Marchenko-Pastur (MP) Principal Component Analysis (PCA) denoising (Veraart et al., 2016a; Veraart et al., 2016b). MP-PCA is designed to denoise *redundant* qMRI data, for which the number of measurements within a given patch of voxels is much larger than the number of linearly-independent contributions to the signal from this patch (i.e., the number of principal components in the low-rank signal representation). MP-PCA applies naturally to acquisition-rich qMRI techniques, such as diffusion MRI, but its application in relaxometry is more challenging due to the limited number of measurements. Here, we exploit synergies between qMRI contrasts enabled by the unified readout, and test whether the combination of less redundant acquisitions with more redundant ones improve MP-PCA denoising of the former.

Our results demonstrate that a unified acquisition enables reliable multi-contrast characterisation of spinal cord microstructure. A unified readout provides metrics that offer complementary information and are intrinsically co-aligned and matched for distortions, without significant losses of resolution for the least redundant modalities. Importantly, data from multiple vendors also demonstrate that MP-PCA is an important pre-processing tool for spinal cord qMRI, with the potential of bringing qMRI one step closer to the clinic by improving the repeatability of microstructural metrics. A unified readout practically increases the redundancy of multi-contrast qMRI data sets, enabling efficient denoising across a range of MRI contrasts.

## 2. Background on MP-PCA

This section outlines key concepts behind MP-PCA denoising required for the understanding of the remainder of the paper.

MP-PCA denoises noisy input matrices **A** = [*a*_*i,j*_] of size *M* × *N* constructed by arranging *M* measurements along rows from *N* neighbouring voxels along columns, such that *M* < *N* without loss of generality. MP-PCA requires *redundant* qMRI protocols, i.e. such that *M* ≫ *P*, with *P* being the number of underlying linearly-independent signal sources. In the absence of noise, *P* identifies the number of non-zero singular values of **A**, which is much smaller than *M* when **A** is redundant. This is the standard assumption behind the PCA-based dimensionality reduction, where one is looking for a P-dimensional hyperplane to represent an M-dimensional measurement. Importantly, in the presence of noise the P-dimensional hyperplane extends in all remaining dimensions, such that **A** becomes full-rank (Veraart et al., 2016a). The denoising problem is then equivalent to identifying the original hyperplane and its dimensionality *P*.

MP-PCA employs random matrix theory to identify information-carrying components within the {*λ*_i_|*i* = 1,…, *M*} singular values of **A**, and to draw a threshold between pure-noise and signal-carrying singular values (Johnstone, 2006). As the pure-noise principal components in the limit *M* ≫ 1 are distributed according to the universal MP distribution, the denoising algorithm identifies the MP distribution within the eigen-spectrum of **A** (Veraart et al., 2016b), and then sets the corresponding *M* − *P* singular values to zero. The *P* information-carrying *above* the right edge of the MP distribution constitute the denoised signal. The output of MP-PCA is 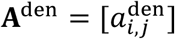, a denoised version of **A**, as well as the simultaneously estimated noise standard deviation σ and the number of signal components (or signal generators) *P*. Note that the estimated *P* will be in general smaller than the original hyperplane dimensionality (without the noise), since some of the informative principal components may fall into the MP noise bulk (Johnstone, 2006). However, we point out that there are no noise-free measurements, and the estimated *P* represents the effective dimensionality of the informative part of the signal, which remains above the noise level. For details of application of random matrix theory and MP-PCA denoising in MRI, please see (Veraart et al., 2016a; Veraart et al., 2016b).

To validate this (or any) denoising method, one can study the distribution *p*(*r*_*i,j*_) of the normalised residuals

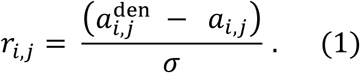

A perfect denoising method removes only Gaussian noise and does not remove tissue anatomy. Hence, as a way to monitor denoising quality, we will check that MP-PCA residuals are Gaussian, i.e.

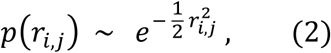

such that the histogram *p*(*r*_*i,j*_) plotted against 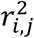 in the semi-log scale is a straight line with slope 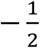 (Veraart et al., 2016b). If tissue anatomy is removed, the histogram plotted as such will not be a straight line, typically blowing up at the tail. Importantly, the line will have a steeper slope if not all noise is removed assuming the no tissue anatomy is spoiled and that σ is correctly estimated, i.e.

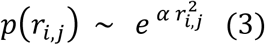

with 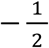 (the removed noise distribution is narrower, and has a smaller variance than the one actually present in the measurement). The performance of denoising increase as the slope *α* approaches 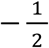 from below (Veraart et al., 2016b), implying that a larger fraction of noise-induced signal variability is mitigated.

## 3 Methods

We synthesised multimodal MRI scans encompassing modalities with different redundancy, emphasising protocols that could be realistically implemented in the spinal cord in vivo, and evaluated the performance of MP-PCA denoising when performed on each modality independent or on multiple modalities jointly.

We also acquired multi-contrast MRI data with unified readout on scanners from two vendors (vendor 1: Philips; vendor 2: Siemens), and characterised the quality of several qMRI metrics as obtained following MP-PCA denoising or without denoising. The shared readout enables the assessment of whether denoising modalities characterised by limited redundancy can be improved if these are denoised jointly with more redundant acquisitions.

In the following sections, we will describe simulations first, as these provide the context for the interpretation of findings in vivo. All analyses were performed using in-house scripts, which are made openly available (http://github.com/fragrussu/PaperScripts/tree/master/sc_unireadout).

### 3.1 In silico study

#### 3.2.1 Signal synthesis

We synthesised realistic spinal cord scans using anatomical information from the Spinal Cord Toolbox (http://github.com/neuropoly/spinalcordtoolbox) (SCT) (De Leener et al., 2017), which contains a high resolution MRI template with voxel-wise volume fractions of white matter (WM, *v*_WM_), grey matter (GM, *v*_GM_) and cerebrospinal fluid (CSF, *v*_CSF_) (Lévy et al., 2015).

Firstly, we used NiftyReg (http://niftyreg.sf.net) `reg_resample` (Modat et al., 2010) with default options to downsample the voxel-wise volume fractions *v*_WM_, *v*_GM_ and *v*_CSF_ to a resolution that is plausible for quantitative MRI of the spinal cord based on EPI (By et al., 2018; Duval et al., 2015; Grussu et al., 2015), i.e. 1×1×5 mm^3^ along R-L, A-P and S-I directions, ensuring realistic partial volume effects. Afterwards, we cropped the field-of-view along the foot-head direction to 200 mm (40 slices), in order to keep a tractable number of synthetic spinal cord voxels to analyse (i.e. 1700 voxels).

We used custom-written Matlab (The MathWorks, Inc., Natick, MA) code to synthesise signals for a rich multimodal quantitative MRI protocol encompassing of DW, qMT, inversion recovery (IR) and multi-echo time (multi-TE) imaging with shared imaging readout (protocol in Table 1, matching our rich in vivo MRI protocol). The total voxel-wise noise-free magnitude signal *S*_TOT_ was obtained as the weighted sum of the signals from WM, GM and CSF, i.e.

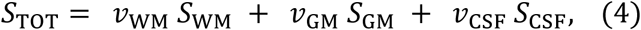

where *v*_WM_ + *v*_GM_ + *v*_CSF_ = 1.

**Table 1.** Sequence parameters used to simulate synthetic multimodal spinal cord scans. In the table, DW, qMT, IR and multi-TE stand respectively for diffusion-weighted, quantitative magnetisation transfer, inversion recovery and multi-echo time. All of DW, qMT, IR and multi-TE imaging rely on the same spin echo EPI readout with long TR (i.e. such that it is hypothesised that TR >> T1). For qMT, each of the 4 repetitions of 11 MT-weighted measurements is characterised by a different delay between the end of the off-resonance pulse train and the readout, i.e. {17, 95, 173, 251} ms. The off-resonance pulse train in qMT was made of 25 sinc-Gaussian pulses (bandwidth: 122 Hz), each lasting 15 ms and with inter-pulse delay of 15 ms (Battiston et al., 2018a).

| Scan | Echo time<br>TE [ms] | Inversion time<br>TI [ms] | Diffusion encoding<br>strength<br>$b$ [s/mm<sup>2</sup>] | Off-resonance pulse<br>flip angle<br>$\theta$ [°] | Off-resonance pulse<br>offset frequency<br>$\Delta f_c$ [KHz] |
| --- | --- | --- | --- | --- | --- |
| **DW**<br>**imaging** | 72 | No inversion<br>pulse used | {0, 300, 1000,<br>2000, 2800} s/mm<sup>2</sup><br>with<br>{8, 4, 10, 18, 28}<br>directions | No off-resonance<br>pulse used | No off-resonance<br>pulse used |
| **qMT**<br>**imaging** | 24 | No inversion<br>pulse used | No diffusion<br>encoding used | 4 repetitions of<br>{0, 426, 433, 524,<br>1429, 1438, 1440,<br>1459, 1460, 1462, 1465} | 4 repetitions of<br>{0.00, 1.07, 1.00, 2.70,<br>14.13, 3.78, 13.60,<br>1.05, 1.01, 3.76, 8.39} |
| **IR**<br>**imaging** | 24 | 12 linearly<br>spaced in<br>[200, 2300] ms | No diffusion<br>encoding used | No off-resonance<br>pulse used | No off-resonance<br>pulse used |
| **multi-TE**<br>**imaging** | {25, 40, 55, 70,<br>85, 100, 200} | No inversion<br>pulse used | No diffusion<br>encoding used | No off-resonance<br>pulse used | No off-resonance<br>pulse used |

For each measurement characterised by sequence parameters (T_E_, T_I_, *b*, ***g***, *θ*, Δ*f*_*c*_) (respectively: echo time, inversion time, diffusion-weighting strength or b-value, diffusion gradient direction, off-resonance pulse flip angle, off-resonance pulse offset frequency), we synthesised each of *S*_WM_, *S*_GM_ and *S*_CSF_ as:

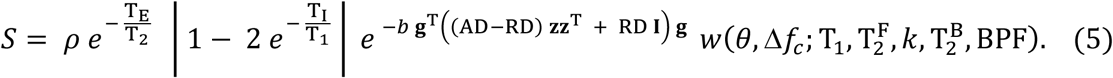

Above, **I** is the 3 × 3 identity matrix, *w* describes MT-weighting, **z** = [0 0 1]^T^ is aligned with the cord longitudinal axis and 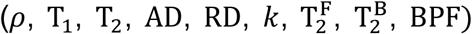 are tissues-specific parameters, in this order: relative proton density (Mezer et al., 2013), macroscopic longitudinal and transverse relaxation rate (Smith et al., 2008), axial and radial diffusivity (Basser et al., 1994), free-to-bound pool exchange rate, free pool transverse relaxation rate, bound pool transverse relaxation rate, bound pool fraction (Henkelman et al., 1993). Eq. 5 models water relaxation as mono-exponential; diffusion as Gaussian, described by an axially symmetric diffusion tensor with primary diffusion direction aligned with the cord longitudinal axis; exchange between free and bound (i.e. myelin) protons according to the two-pool MT model (Henkelman et al., 1993). The MT-weighting factor **z** was calculated via direct numerical integration of the two-pool Bloch equations (details in Supplementary Material S1), assuming a super-Lorentzian line shape for bound protons and simulating off-resonance pulse trains made of 25 sinc-Gaussian pulses (bandwidth: 122 Hz), each lasting 15 ms and with inter-pulse delay of 15 ms, as used before in spinal cord application (Battiston et al., 2018a).

We synthesised a unique noise-free signal profile in each tissue voxel by simulating within-tissue variability in WM and GM. This ensures that each synthetic voxel has its own unique sources of signal, avoiding obvious redundancies within the set of synthetic signals, as these could lead to overestimation of the performances of MP-PCA denoising (Ades-Aron et al., 2018). In practice, we drew voxel-wise values for each of 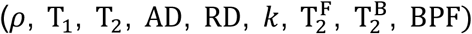 from a tissue-specific Gaussian distribution, with parameters inspired by values known from literature (Battiston et al., 2018a; Grussu et al., 2015; Smith et al., 2008) (parameters in Table 2).

**Table 2.** Tissue parameters used to generate the synthetic spinal cord scans. Values are inspired by previous literature (Battiston et al., 2018a; Grussu et al., 2015; Smith et al., 2008). For white/grey matter, within-tissue variability was simulated by drawing parameter values from a Gaussian distribution and assigning the obtained values to different voxels. The mean and standard deviation of the Gaussian distributions are reported in the table (standard deviation within brackets, equal to 10% of the mean). For the cerebrospinal fluid (CSF), tissue parameters were fixed to the same values across all CSF-containing voxels.

| **Tissue** | Relative proton density<br>$\rho$ | Longitudinal relaxation time<br>$T_1$ [ms] | Transverse relaxation time<br>$T_2$ [ms] | Axial diffusivity AD<br>[$\mu\text{m}^2/\text{ms}$] | Radial diffusivity RD<br>[$\mu\text{m}^2/\text{ms}$] | Free-to-bound proton exchange rate<br>$k$ [1/s] | Free proton transverse relaxation time $T_2^F$ [ms] | Bound proton transverse relaxation time $T_2^B$ [$\mu\text{s}$] | Bound pool fraction BPF |
| --- | --- | --- | --- | --- | --- | --- | --- | --- | --- |
| **White matter** | 0.70<br>(0.07) | 1000<br>(100) | 70<br>(7) | 2.10<br>(0.21) | 0.40<br>(0.040) | 2.3<br>(0.23) | We fix<br>$T_2^F = T_2$ | 12<br>(0.12) | 0.14<br>(0.014) |
| **Grey matter** | 0.80<br>(0.08) | 1200<br>(120) | 80<br>(8) | 1.60<br>(0.16) | 0.55<br>(0.055) | 1.7<br>(0.17) | We fix<br>$T_2^F = T_2$ | 12<br>(0.12) | 0.08<br>(0.008) |
| **CSF** | 1.00 | 4000 | 800 | 3.00 | 3.00 | Not defined<br>(BPF = 0) | We fix<br>$T_2^F = T_2$ | Not defined<br>(BPF = 0) | 0.00 |

The synthetic spinal cord phantom is made openly available online (http://github.com/fragrussu/PaperScripts/tree/master/sc_unireadout/sc_phantom).

#### 3.2.2 Denoising

We corrupted the synthetic signals with Gaussian noise at different signal-to-noise ratios (SNRs) (300 unique noise realisations on 1700 voxels), ranging from 10 to 40 (SNR evaluated with respect to the *b* = 0 signal in WM for the DW measurements).

Afterwards, we used the Matlab implementation of the MP-PCA algorithm (http://github.com/NYU-DiffusionMRI/mppca_denoise) to denoise the synthetic spinal cord images at the various SNRs.

For our simulations, we processed MP-PCA matrices constructed by arranging spinal cord voxels within an individual MRI slice along rows and different MRI measurements along columns (i.e. slice-by-slice cord denoising). We implemented three different denoising strategies:

1. individual denoising of each modality among DW, IR, multi-TE and quantitative MT imaging respectively (importantly, IR, multi-TE have limited redundancy and would not theoretically qualify for MP-PCA, which is expected to remove little to no noise);
2. joint denoising of all modalities concatenated as one large set of measurements;
3. joint denoising of DW imaging concatenated with each of IR, multi-TE and qMT imaging in series respectively, which would be useful to describe cases when only one modality other than DW imaging is acquired.

#### 3.2.3 Analysis

We evaluated the performance of MP-PCA denoising by studying the percentage relative error *ɛ* between the denoised signals *S*_TOT,denoised_ and the ground truth signal *S*_TOT_. We estimated accuracy and precision of the different denoising strategies by calculating respectively the median of *ɛ* (such that the closer to zero, the higher the accuracy) and interquartile range (IQR) of *ɛ* (such that the lower, the higher the precision) within the synthetic spinal cord over the 100 noise instantiations. We also studied the distributions of normalised residuals *p*(*r*), evaluated according to Eq. 1.

### 3.3 In vivo study

We performed clinically viable, multi-contrast spinal cord qMRI scans on healthy volunteers and analysed them to characterise the performance of MP-PCA denoising on different qMRI modalities, devising denoising strategies for acquisitions with different levels of redundancy. The experimental sessions were approved by local research ethics committees.

Our qMRI protocols exhibit a unified signal readout, which is based on spin echo EPI, a typical choice for DW imaging. The shared readout ensures comparable noise characteristics across the different qMRI modalities, thus enabling joint denoising of different qMRI contrasts. The MRI protocol in vendor 1 encompasses DW, qMT, IR and multi-TE imaging, while in vendor 2 includes DW and multi-TE imaging. The MRI protocol in vendor 2 is less rich due to practical availability of pulse sequences. Nonetheless, it suffices to demonstrate the potential of joint multi-contrast denoising of modalities with different redundancies, and is representative of protocols required in multi-contrast techniques such as TEDDI (Veraart et al., 2018).

In all systems, MRI scans were performed axially-oblique at the level of the cervical cord, with filed-of-view centred at the C2-C3 intervertebral disk (foot-head coverage of 60 mm).

#### 3.3.1 MRI: vendor 1

The protocol developed on a 3T Philips Achieva machine, located at the UCL Queen Square Institute of Neurology (London, UK) consisted of multi-contrast, single-shot spin echo EPI scans with unified signal readout based on reduced field-of-view ZOOM technology (Wheeler‐Kingshott et al., 2002), which enable 4 contrast mechanisms to be exploited: DW imaging, qMT imaging, IR imaging and multi-TE imaging (mTE, i.e. acquisitions of single-shot images at different TE). Salient sequence parameters, including information on b-values, echo/inversion times, off resonance saturation and cardiac gating are reported in Table 3.

**Table 3.** Salient sequence parameters for the qMRI protocol with unified readout implemented on vendor 1 (Philips Achieva, London, UK). DW, qMT, IR and multi-TE stand for diffusion-weighted, quantitative magnetisation transfer, inversion recovery and multi-echo time. Consistently with simulations, in qMT each repetition is characterised by a different delay between the end of the off-resonance train and the readout, i.e. {17, 95, 173, 251} ms. qMT off-resonance trains were made of 25 sinc-Gaussian pulses (bandwidth: 122 Hz), each lasting 15 ms and with inter-pulse delay of 15 ms (Battiston et al., 2018).

| Scan | TE [ms] | TR [ms] | RL-AP field-of-view [mm<sup>2</sup>] | Resolution [mm<sup>2</sup>] | Slices | Bandwidth [Hz/pixel] | Acceleration | Parameters for quantitative imaging | Nominal scan time |
| --- | --- | --- | --- | --- | --- | --- | --- | --- | --- |
| **DW imaging** | 71 | 12000 (peripheral gating, delay of 150 ms) | 64x48 | 1x1 | 12 slices, 5 mm-thick, 3 packages | 2132 | Half-scan factor of 0.6 | 8 b = 0; b = {300, 1000, 2000, 2800} s mm<sup>-2</sup> with {4, 10, 18, 28} gradient directions | 16 min : 41 s |
| **qMT imaging** | 24 | 7246 | 64x48 | 1x1 | 12 slices, 5 mm-thick, 3 packages | 2132 | Half-scan factor of 0.6 | 1 non-MT weighted, 10 MT-weighted; 4 repetitions with different slice ordering (same off-resonance pulses as in Table 1) | 16 min : 18 s |
| **IR imaging** | 24 | 8305 | 64x48 | 1x1 | 12 slices, 5 mm-thick, 3 packages | 2132 | Half-scan factor of 0.6 | 12 linearly-spaced inversion times TI in [100; 2300] ms; spacing of 200 ms | 4 min : 50 s |
| **multi-TE imaging** | Various | 12000 (1 dummy scan per TE) | 64x48 | 1x1 | 12 slices, 5 mm-thick, 3 packages | 2132 | Half-scan factor of 0.6 | 7 echo times TE: {25, 40, 55, 70, 85, 100, 200} ms | 2 min : 48 s |

The protocol also included an anatomical 3D FFE scan (flip angle of 7°, TE of 4.1 ms, TR of 20 ms, resolution of 0.75 × 0.75 × 5 mm^3^ and field-of-view of 180 × 240 × 60 mm^3^ along R-L, A-P, S-I directions; ProSet fat suppression, 3 signal averages, scan time of 3 min: 30 s) and standard B0 and B1 field mapping for accurate qMT analysis. Both B0 and B1 mapping were based on 3D FFE acquisitions with resolution of 2.25 × 2.25 × 5 mm^3^ and field-of-view of 215 × 206 × 60 mm^3^ along R-L, A-P, S-I directions. B0 mapping was performed with the double-echo method (Jezzard and Balaban, 1995), with parameters: flip angle of 25°, TE of 6.9 ms and 9.2 ms, TR of 50 ms, scan time of 1 min: 40 s. B1 mapping was instead performed via actual flip angle imaging (Yarnykh, 2007), with parameters: flip angle of 60°, TE of 2.5 ms, TR of 30 ms, TR extension of 120 ms, scan time of 1 min: 40 s).

The nominal acquisition time was roughly 47 min, with variations depending on subject’s heart rate. We scanned 4 healthy volunteers twice (2 males, age range 28-40), with the rescan performed within one month of the first scan.

#### 3.3.2 MRI: vendor 2

For vendor 2, we performed scans on two separate 3T Siemens Prisma systems, located at the New York University School of Medicine (USA) and at the Neuroimaging Functional Unit of the University of Montreal (Canada).

The protocol consisted in exploiting 2 contrast mechanisms including DW imaging and multi-TE imaging with unified readout based on syngo ZOOMit reduced field-of-view technology (Rieseberg et al., 2002) (salient parameters including b-values, TEs and cardiac gating are reported in Table 4). The protocol also included a 3D MEDIC scan for anatomical depiction (flip angle of 30°, TE of 15 ms, TR of 625 ms, resolution of 0.50 × 0.50 × 5 mm^3^ and field-of-view of 128 × 128 × 60 mm^3^ along R-L, A-P, S-I directions; 3 signal averages, scan time of 6 min: 24 s).

**Table 4.** Salient sequence parameters for the qMRI protocol with unified readout implemented on vendor 2 (Siemens Prisma systems in New York, USA and Montreal, Canada). DW and and multi-TE stand respectively for diffusion-weighted and multi-echo time.

| Scan | TE<br>[ms] | TR<br>[ms] | RL-AP<br>field-of-view<br>[mm<sup>2</sup>] | Resolution<br>[mm<sup>2</sup>] | Slices | Bandwidth<br>[Hz/pixel] | Acceleration | Parameters for<br>quantitative<br>imaging | Nominal<br>scan<br>time |
| --- | --- | --- | --- | --- | --- | --- | --- | --- | --- |
| **DW**<br>**imaging** | 85 | 9194<br>(peripheral<br>gating,<br>delay of<br>200 ms) | 64×64 | 1×1 | 12 slices,<br>5 mm-thick,<br>12<br>concatenations | 1240 | 6/8 Partial<br>Fourier<br>Imaging | 4 b = 0;<br>b = {300, 1000,<br>2000, 2800}<br>s mm<sup>-2</sup> with<br>{6, 12, 20, 30}<br>directions in NY<br>and<br>{20, 20, 20, 30}<br>directions in<br>Montreal | 11 min : 02 s<br>in NY;<br>14 min : 24 s<br>in Montreal<br>(a higher<br>number of<br>DW images<br>acquired in<br>Montreal) |
| **multi-TE**<br>**imaging** | various | 13000<br>(peripheral<br>gating,<br>delay of<br>200 ms) | 64×64 | 1×1 | 12 slices,<br>5 mm-thick,<br>12<br>concatenations | 1240 | 6/8 Partial<br>Fourier<br>Imaging | 7 echo times<br>TE:<br>{45, 60, 75, 90,<br>105, 120, 200}<br>ms | 1 min : 31 s |

The total scan time was 18 min: 57 s in the New York Prisma and 22 min: 19 s in the Montreal Prisma, with the scan time difference due to slightly higher number of diffusion directions being acquired in Montreal. Two subjects were scanned in New York (1 male, 28 years old; 1 female, 25 years old) and one subject (male, 28 years old) in Montreal after obtaining informed written consent. The vendor-provided 64 channel head-neck coil was used in both cases for signal reception.

#### 3.3.3 Denoising

We implemented the same denoising strategies as in simulations:

1. individual denoising of each modality separately;
2. joint denoising of all modalities together;
3. joint denoising of DW imaging concatenated with each of IR, multi-TE and qMT imaging in series (multi-TE only for Prisma).

We performed denoising slice-by-slice to account for the anisotropic voxel-size and to limit the effect of potential between-shot signal fluctuations due to physiological noise (Summers et al., 2006). We proceeded as follows:

- the spinal cord was identified on the mean DW image with SCT sct_propseg (De Leener et al., 2014);
- all cord voxels of an MRI slice were arranged as one matrix and denoised with MP-PCA;
- noise floor (Gudbjartsson and Patz, 1995) was subsequently mitigated on the denoised signals with the method of moments (Koay and Basser, 2006).

#### 3.3.4 Post-processing

We performed motion correction on the concatenation of all acquired EPI images within an MRI session. Practically, we ran slice-wise rigid motion correction with **sct_dmri_moco** on the non-denoised scans, treating qMT, IR and multi-TE images as *b* = 0 scans. The estimated registration transformations were stored and used to correct all the denoised versions of each qMRI modality, as well as the non-denoised data. This was done to focus our analysis on the effect that thermal noise removal has on qMRI metrics.

Afterwards, we segmented the whole cord and the grey matter in the anatomical spinal cord scan respectively with **sct_propseg** and with **sct_deepseg_gm**. We also segmented the spinal cord in the mean DW EPI image with **sct_propseg**.

Lastly, we co-registered the anatomical spinal cord scan to the mean EPI image with **sct_register_multimodal**, using dilated spinal cord masks in the two image spaces to guide registration (dilation performed with NifTK **seg_maths**, available at http://github.com/NifTK/NifTK). The estimated warping field transformation was used to warp the grey matter mask to EPI space, which was subsequently used to obtain a white matter mask by subtracting it from the whole-cord mask. For vendor 1, the warping field was also used to resample the B0 and B1 magnetic field maps to the EPI space for downstream model fitting.

#### 3.3.5 Evaluation of quantitative metrics

We fit quantitative models/signal representations for the different contrasts and obtain popular metrics that are promising imaging biomarkers. These were:

- diffusion kurtosis imaging (Jensen et al., 2005; Veraart et al., 2011) on DW data (both vendors) with DiPy **dipy.reconst.dkimodule** (http://nipy.org/dipy/examples_built/reconst_dki.html), obtaining voxel-wise diffusion and kurtosis tensors, of which fractional anisotropy (FA), mean diffusivity (MD) and mean kurtosis (MK) were considered for downstream analyses;
- mono-exponential T_2_ relaxation on multi-TE data (both vendors) with MyRelax **getT2T2star.py** (http://github.com/fragrussu/myrelax), obtaining voxel-wise macroscopic T_2_;
- mono-exponential T_1_ relaxation on IR data (vendor 1 only), with MyRelax getT1IR.py, obtaining voxel-wise macroscopic T_1_;
- two-pool qMT model on qMT data (vendor 1 only) with custom-written Matlab code (Battiston et al., 2018a), obtaining voxel-wise bound pool fraction BPF, exchange rate *k* and bound pool transverse relaxation 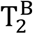, of which BPF and *k* were considered for downstream analyses. For qMT fitting, static/transmitted fields were corrected on a voxel-by-voxel basis using the B0 and B1 field maps warped to EPI space.

#### 3.3.6 Analysis

We calculated the normalised signal residuals (Eq. 1) for all denoising approaches and for all subjects, sessions and vendors, and evaluated their distributions within the spinal cord.

We also characterised values of all qMRI metrics by calculating the median within grey and white matter for all denoising strategies (including no denoising).

Finally, we quantified each metric variability by calculating a percentage coefficient of variation (CoV) within grey matter and within white matter for all denoising strategies (including no denoising). We defined CoV as

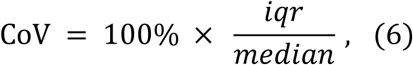

where *iqr* is the interquartile range of a metric within grey/white matter, measuring the metric variability, while *median* is the median value of the metric within the same tissue. We hypothesise that effective denoising would reduce noise-induced metric variability, resulting in lower *iqr* and unchanged *median* and hence lower CoV, under assumption that variability due to noise is much larger than the biological variability (please see figure 4 of (Ades-Aron et al., 2018)).

## 4 Results

### 4.1 In silico study

Fig. 1 shows distributions of residuals (in the semilog scale, plotted against the squared residuals) from simulations. These are reported for the different qMRI modalities and for different denoising strategies. In such plots, Gaussian residuals align along a straight line. The charts reveal that residuals are Gaussian for at least 3*σ*. Moreover, they also highlight that joint multi-contrast MP-PCA denoising mitigates more efficiently noise for those modalities featuring limited redundancy than if denoised independently, such as IR imaging and mTE imaging: their residuals are closer to the ideal unit Gaussian when denoised jointly with more redundant modalities than when denoised alone. In contrast, denoising performance does not improve for joint multi-contrast MP-PCA of those modalities that are intrinsically highly redundant (i.e. DW and qMT imaging).

**Fig. 1.**
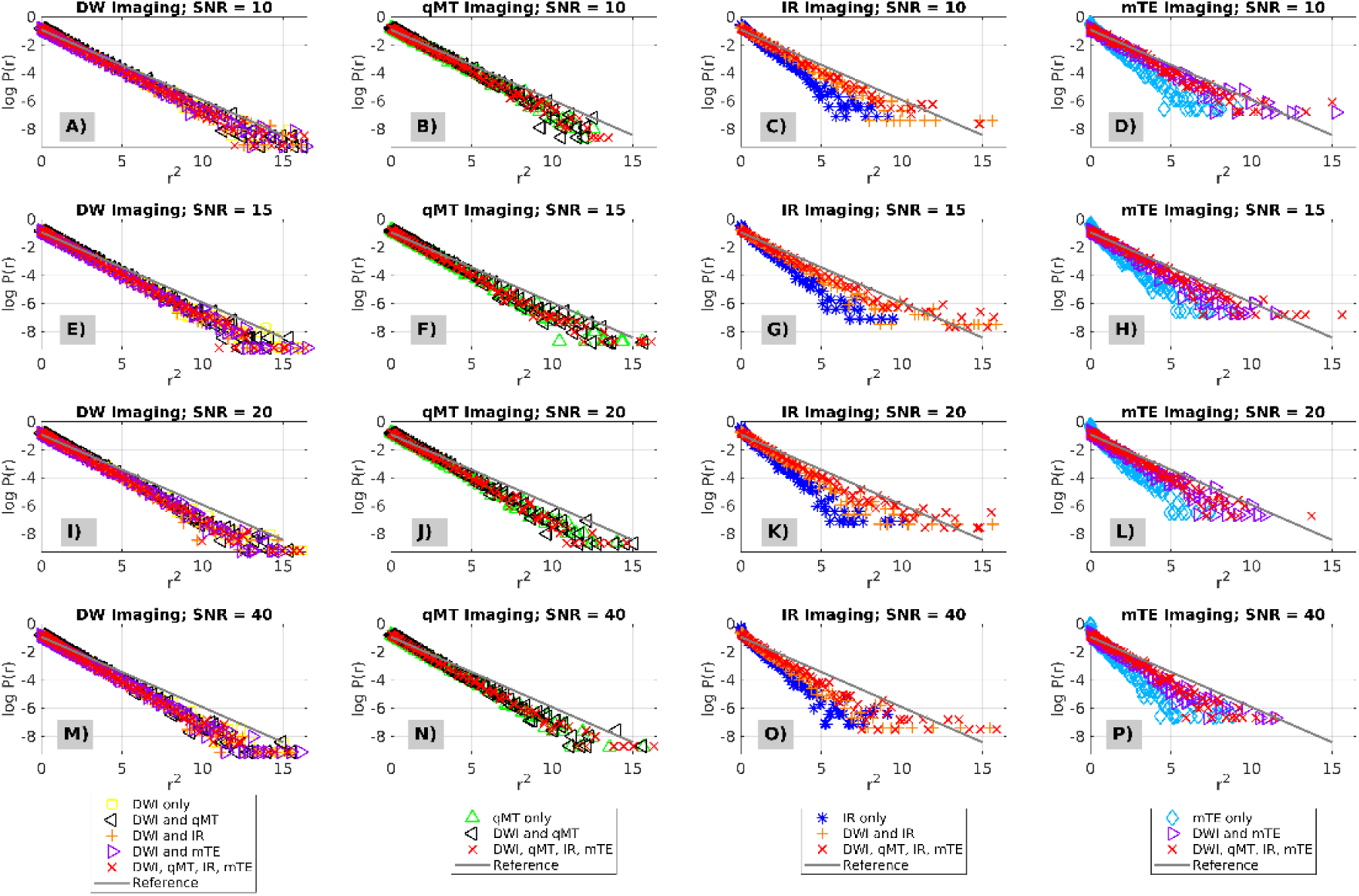
Distributions of normalised residuals from simulations for different denoising strategies. Each plot reports the logarithm of the residual histogram, scattered against the corresponding values of squared residuals. In these plots, Gaussian residuals appear as straight lines. First row (panels A to D): simulated SNR of 10; second row (panels E to H): simulated SNR of 15; third row (panels I to L): simulated SNR of 20; fourth row (panels M to P): simulated SNR of 40. SNR is evaluated with respect to the synthetic white matter signal in the diffusion scan at *b* = 0. Different columns refer to different MRI scans: from left to right, DW imaging (panels A, E, I, M), qMT imaging (panels B, F, J, N), IR imaging (panels C, G, K, O), mTE imaging (panels D, H, L, P).

Fig.2 shows percentage relative error accuracy (top row: error median) and precision (bottom row: error IQR) of the denoised signals compared to the noise-free ground truth, for different qMRI modalities and different denoising strategies. While plots do not highlight any noticeable differences in terms of accuracy for the different denoising strategies (i.e. joint denoising or individual denoising, since their confidence intervals overlap perfectly), they do suggest that better precision (i.e. error IQR closer to zero) can be achieved for modalities that are intrinsically limited in redundancy, when these are denoised jointly with more redundant modalities. For example, IQR drops from 8% to 5% at SNR = 10 for mTE when it is denoised jointly with all modalities, as compared to when mTE is denoised along. Similar to what is reported in Fig. 1, no appreciable improvement of denoising performance is observed with joint multi-contrast denoising for modalities that intrinsically feature high redundancy. This is apparent for qMT and even more so for DW imaging, since their percentage error IQR does not change when these are denoised jointly with other modalites).

**Fig. 2.**
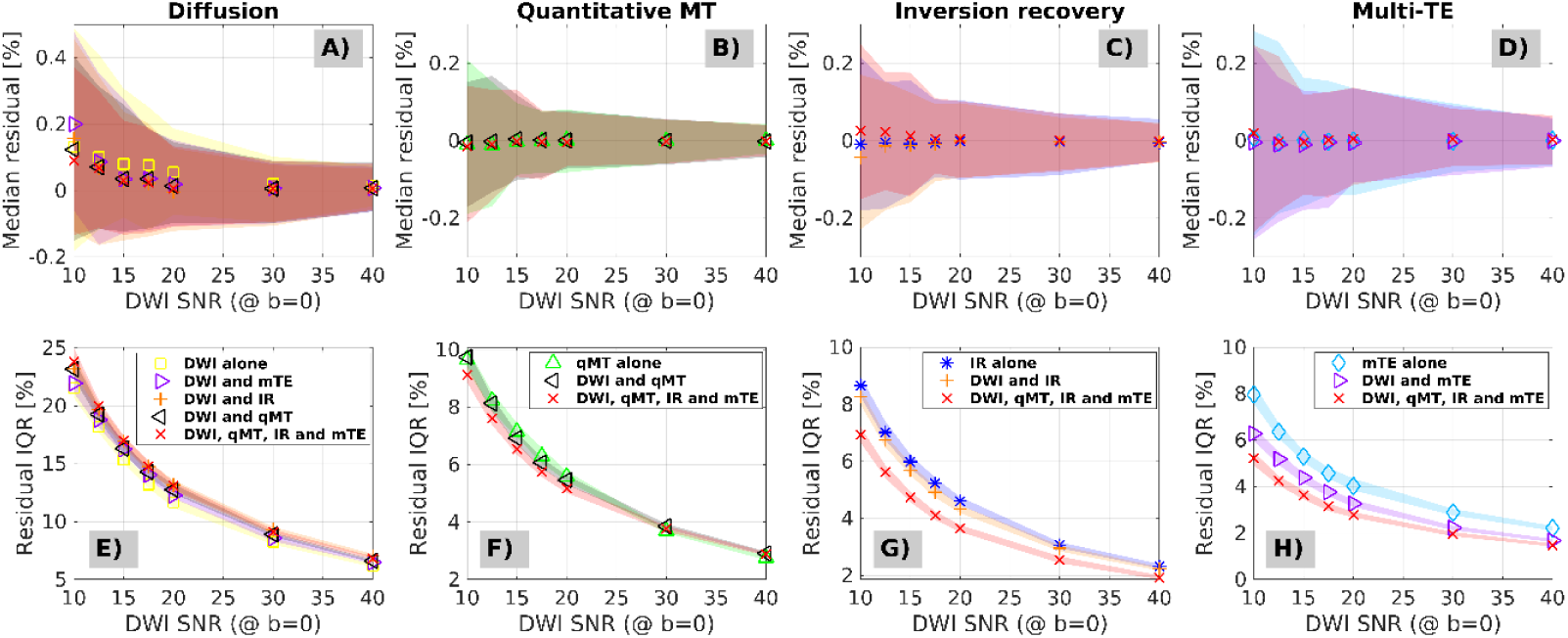
Accuracy and precision of different denoising strategies as obtained from percentage relative errors (percentage errors between denoised signals and noise-free ground truth signals) in simulations. Panels A to D (top) show median percentage at different SNR levels, and represent a measure of accuracy (the closer to zero, the higher the accuracy; DW imaging in A, qMT imaging in B, IR imaging in C, mTE imaging in D). Panels E to H (bottom) show percentage relative error interquartile ranges at different SNR levels, and represent a measure of precision (the closer to zero, the higher the precision; DW imaging in E, qMT imaging in F, IR imaging in G, mTE imaging in H).

### 4.2 In vivo study

Figs. 3 and 4 show examples of acquired and denoised in vivo images alongside distributions of residuals. Fig. 3 illustrates information for vendor 1, while Fig. 4 for vendor 2. For both vendors, clear improvements in image quality are visually and quantitatively observed, especially for DW imaging. Residuals are Gaussian for at least 3*σ*, and multimodal denoising mitigates noise more efficiently for those modalities that exhibit limited intrinsic redundancy (i.e. IR and mTE imaging) as compared to individual denoising (where residuals are further away from the ideal unit Gaussian).

**Fig. 3.**
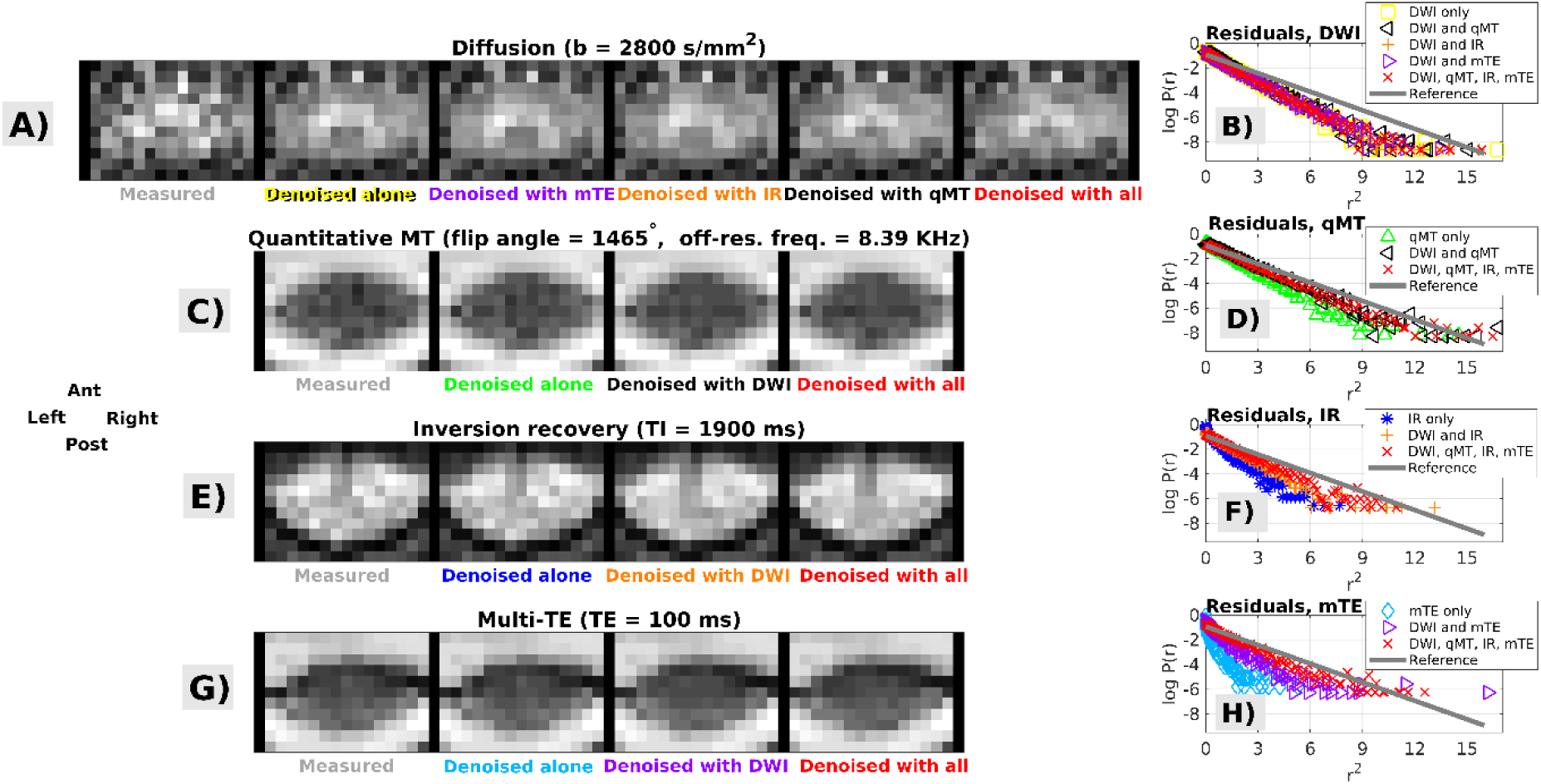
Examples of MP-PCA denoising in one subjects who was scanned with vendor 1. Panels A, C, E, G show raw and denoised images, obtained according to different strategies; panels B, D, F, H show the corresponding distributions of normalised residuals in log-scale, plotted against the squared residuals (in these plots, Gaussian residuals align along a straight line). DW imaging: panels A (images) and B (residuals); qMT imaging: panels C (images) and D (residuals); IR imaging: panels E (images) and F (residuals); mTE imaging: panels G (images) and H (residuals). *Ant*, *Post*, *Right*, *Left* respectively indicate subject’s anterior, posterior parts and right and left sides.

**Fig. 4.**
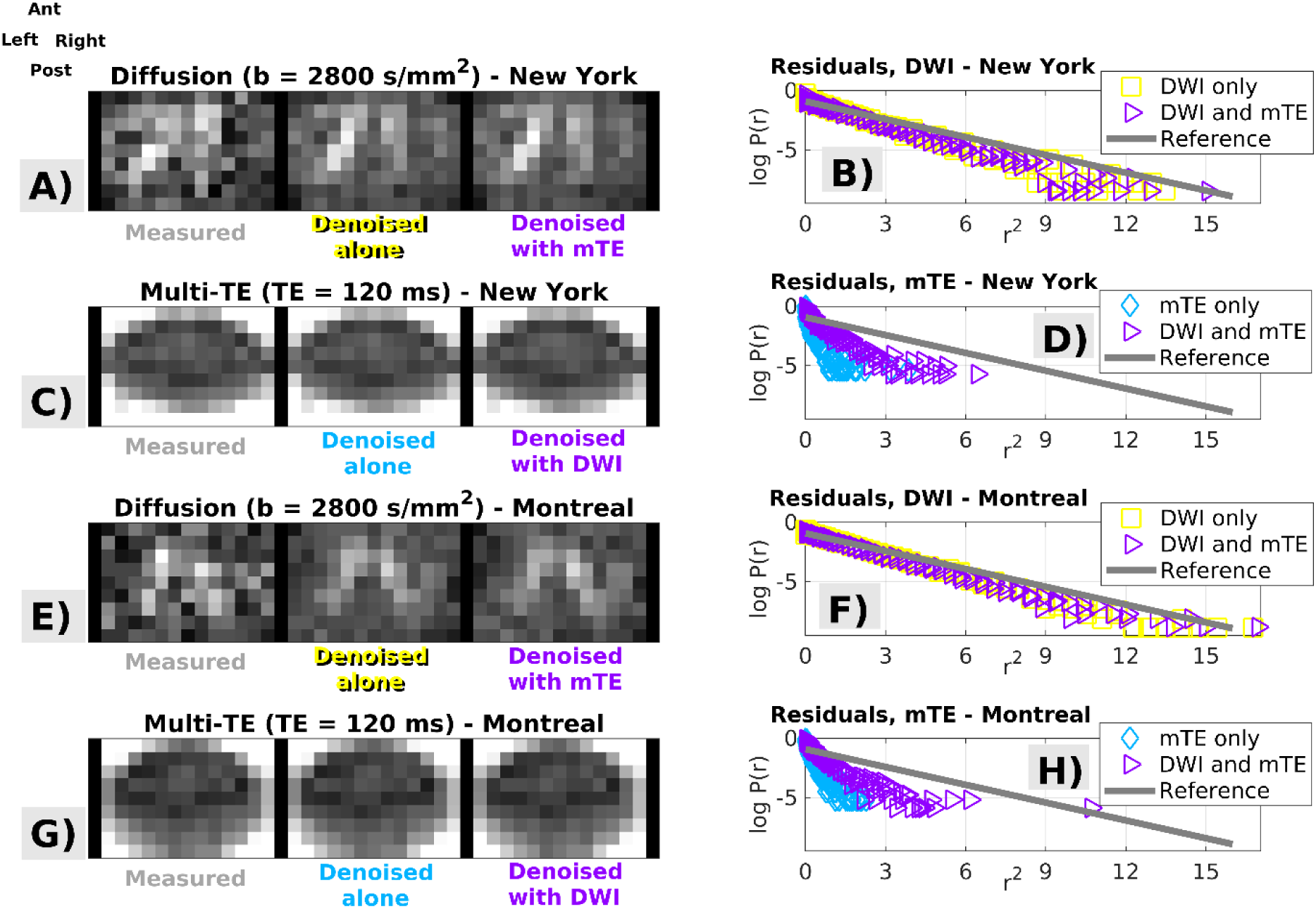
Examples of MP-PCA denoising in two subjects, scanned with vendor 2 respectively in New York and in Montreal. Panels A, C, E, G show raw and denoised images, obtained according to different strategies; panels B, D, F, H show the corresponding distributions of normalised residuals in log-scale, plotted against the squared residuals (in these plots, Gaussian residuals align along a straight line). DW imaging: images in panels A and E, residuals in B and F; mTE imaging: images in panels C and G, residuals in panels D and H. *Ant*, *Post*, *Right*, *Left* respectively indicate subject’s anterior, posterior parts and right and left sides.

Figs. 5, 6 and 7 show examples of quantitative parametric maps obtained in one representative subject from vendor 1 (Fig. 5; DW, qMT, IR and mTE imaging) and from the vendor 2 (New York system in Fig. 6, Montreal system in Fig. 7; DW and mTE imaging) for different denoising strategies. Visual inspection suggests that MP-PCA denoising generates less noisy maps, especially for vendor 1. The most striking examples of improved parameter estimation are seen in both vendors for DW imaging parameter MK. Additionally, improvements on visual inspection are apparent for other qMRI metrics such as BPF and T_1_, especially for joint multimodal denoising.

**Fig. 5.**
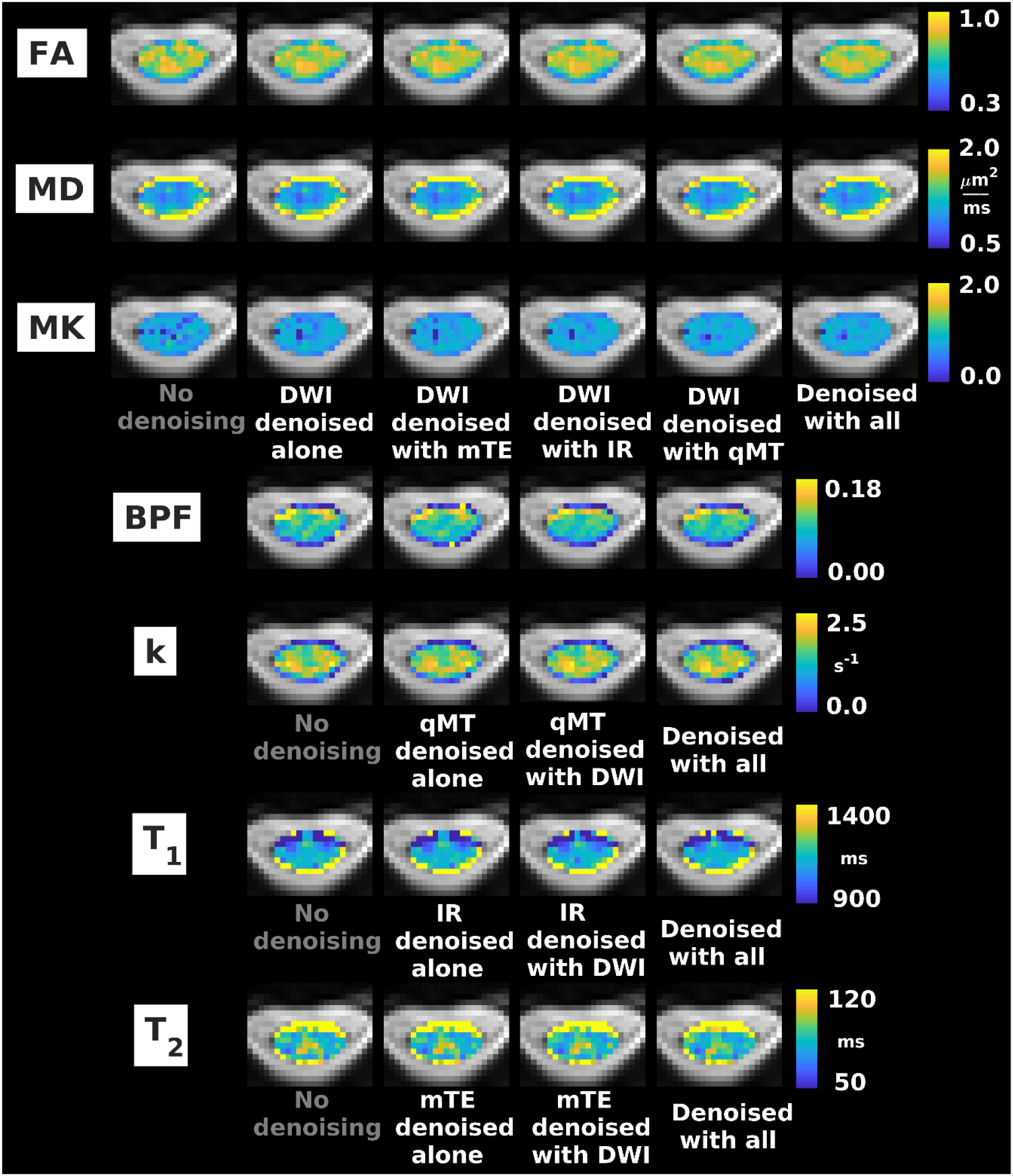
Examples of quantitative maps from vendor 1. From top to bottom: FA, MD, MK (DW imaging); BPF, k (qMT imaging); T1 (IR imaging); T2 (mTE imaging). Different rows illustrate the metrics obtained according to different denoising strategies (no denoising; independent denoising of each modality; various combinations of joint multi-modal denoising). Quantitative maps are overlaid onto the mean non-DW image and shown within the cord only. The same anatomical conventions with regard to subject’s anterior, posterior parts and right and left sides as in Fig. 3 are used.

**Fig. 6.**
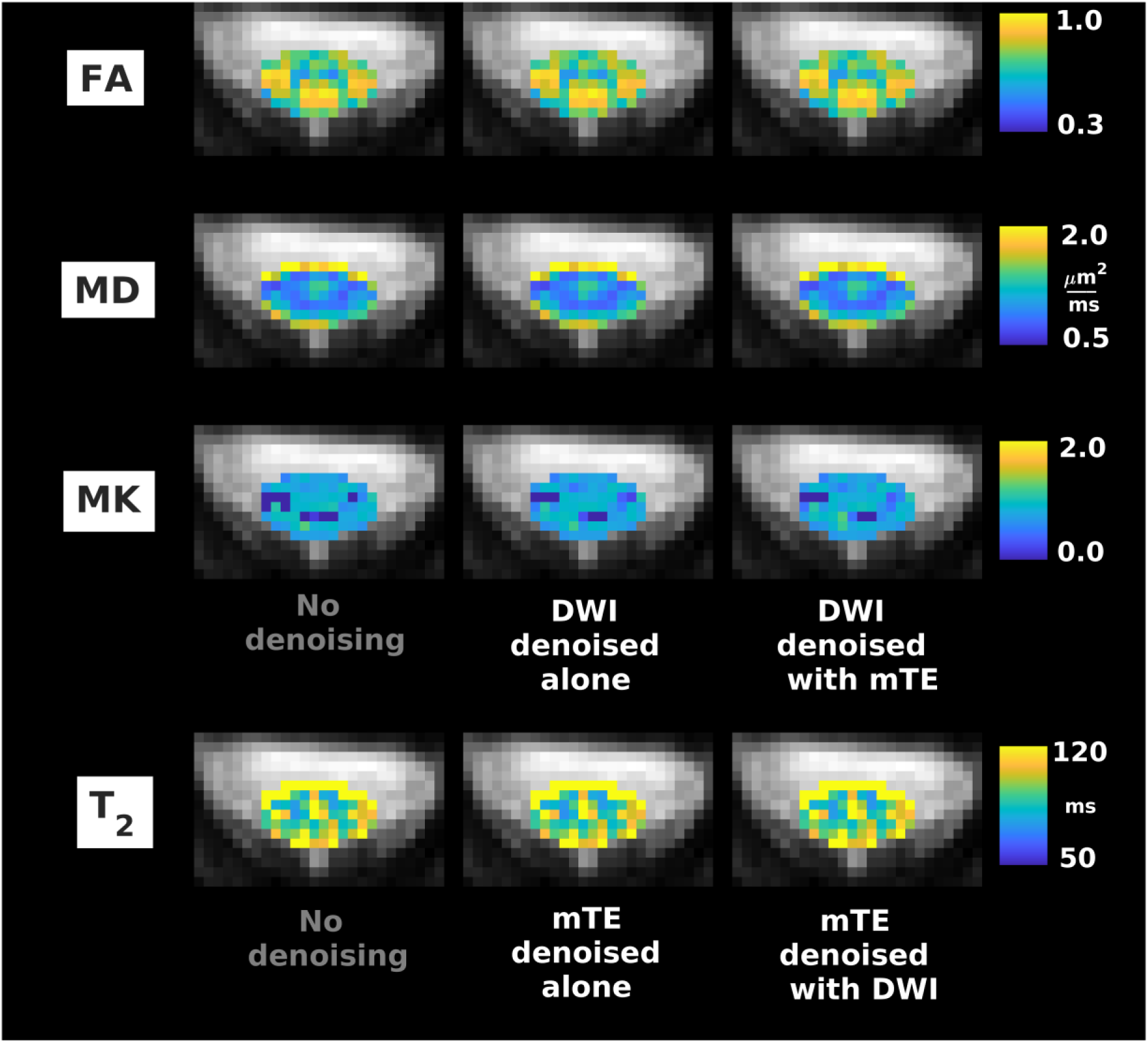
Examples of quantitative maps from vendor 2 (Siemens Prisma system located in New York, USA). From top to bottom: FA, MD, MK (DW imaging); T2 (mTE). Different rows illustrate the metrics obtained according to different denoising strategies (no denoising; independent denoising of each modality; various combinations of joint multi-modal denoising). Quantitative maps are overlaid onto the mean non-DW image and shown within the cord only. The same anatomical conventions with regard to subject’s anterior, posterior parts and right and left sides as in Fig. 4 are used.

**Fig. 7.**
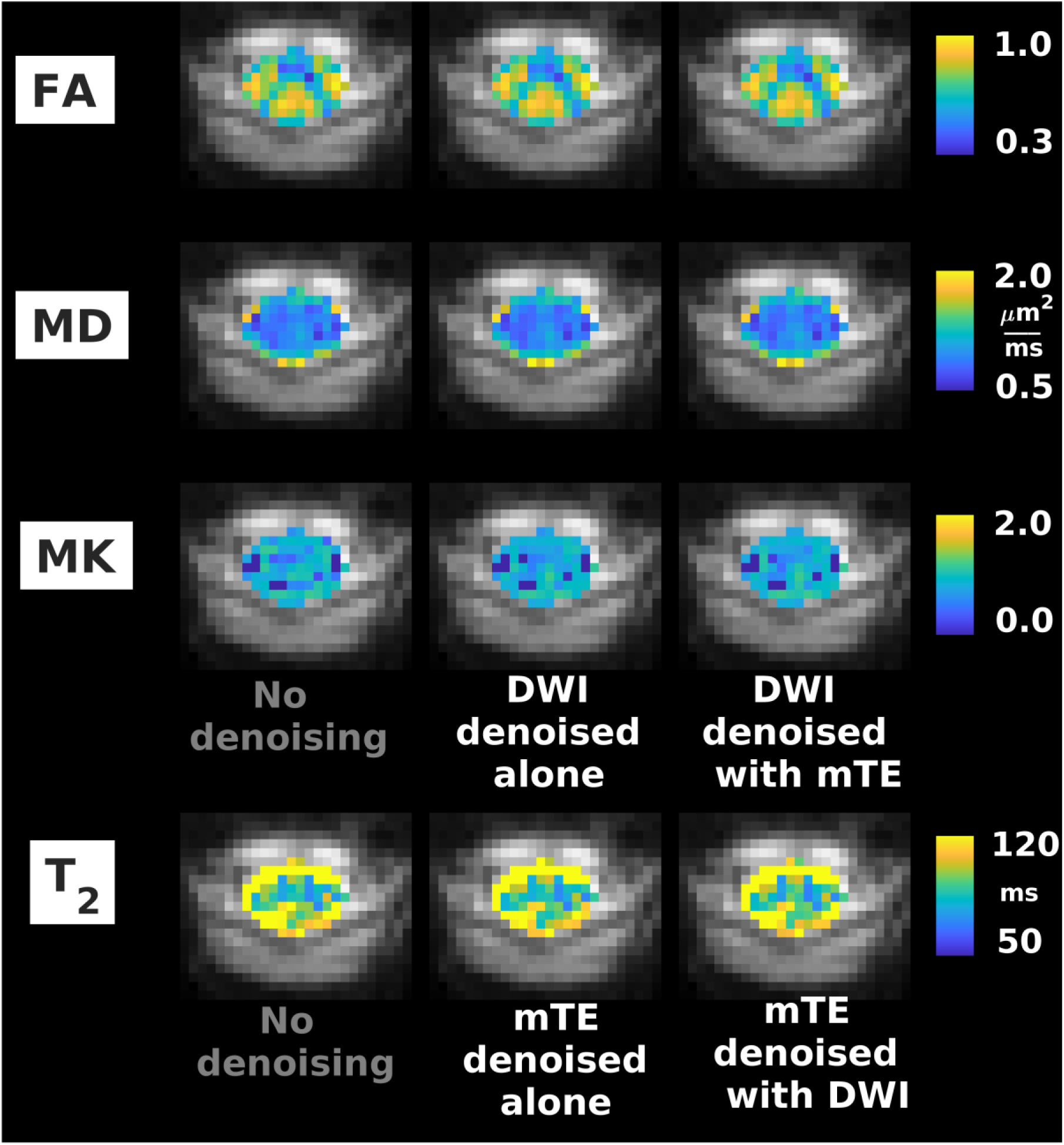
Examples of quantitative maps from vendor 2 (Siemens Prisma system located in Montreal, Canada). From top to bottom: FA, MD, MK (DWI); T2 (mTE). Different rows illustrate the metrics obtained according to different denoising strategies (no denoising; independent denoising of each modality; various combinations of joint multi-modal denoising). Quantitative maps are overlaid onto the mean non-DW image and shown within the cord only. The same anatomical conventions with regard to subject’s anterior, posterior parts and right and left sides as in Fig. 4 are used.

Tables 5 and 6 report median values of all qMRI metrics in grey and white matter for the different denoising strategies (Table 5: vendor 1; Table 6: vendor 2, pooling together results from the two systems). The tables reveal contrasts between grey and white matter in various metrics. Examples that are consistent between vendors include: higher FA and MD in white compared to grey matter; similar MK in grey/white matter; slightly higher T_2_ in white compared to grey matter. Other examples from vendor 1 include: similar BPF and T_1_ in grey/white matter; higher exchange rate *k* in grey compared to white matter. The tables also show that systematic differences between the data sets acquired with the two vendors exist, as for example: higher T_2_ and MK and lower MD in data from vendor 2 compared to 1; different grey/white matter contrasts in FA. Notably, Tables 5 and 6 also demonstrate that denoising introduce little to no biases in the quantitative parametric maps. In all cases and for both vendors the tissue-wise medians never differ for more than 5% compared to the values obtained without any denoising.

**Table 5.** Median values in grey and white matter of qMRI metrics obtained with a scanner from vendor 1 following different denoising strategies. Median values from different subjects and scans are pooled so that figures in the table report mean and standard deviation (in brackets) across subjects and scans. In all cases, values of metrics obtained after denoising are less than 5% different from the values obtained with no denoising.

|  | **No**<br>**denoising** | **Individual**<br>**denoising** | **Joint**<br>**denoising**<br>**DWI-mTE** | **Joint**<br>**denoising**<br>**DWI-IR** | **Joint**<br>**denoising**<br>**DWI-qMT** | **Joint**<br>**denoising**<br>**of all** |
| --- | --- | --- | --- | --- | --- | --- |
| **FA**<br>**(DWI)** | GM: 0.70 (0.05)<br>WM:0.73 (0.04) | GM: 0.70 (0.06)<br>WM:0.73 (0.04) | GM: 0.70 (0.06)<br>WM:0.74 (0.04) | GM: 0.70 (0.06)<br>WM:0.73 (0.04) | GM: 0.71 (0.06)<br>WM:0.74 (0.04) | GM: 0.71 (0.06)<br>WM:0.73 (0.04) |
| **MD [$\mu\text{m}^2/\text{ms}$]**<br>**(DWI)** | GM: 1.01 (0.08)<br>WM:1.24 (0.09) | GM: 1.02 (0.08)<br>WM:1.24 (0.09) | GM: 1.02 (0.08)<br>WM:1.24 (0.10) | GM: 1.02 (0.08)<br>WM:1.24 (0.10) | GM: 1.03 (0.08)<br>WM:1.24 (0.10) | GM: 1.03 (0.08)<br>WM:1.24 (0.10) |
| **MK**<br>**(DWI)** | GM: 0.75 (0.18)<br>WM:0.75 (0.13) | GM: 0.75 (0.18)<br>WM:0.76 (0.11) | GM: 0.75 (0.20)<br>WM:0.77 (0.11) | GM: 0.74 (0.21)<br>WM:0.76 (0.11) | GM: 0.76 (0.19)<br>WM:0.78 (0.11) | GM: 0.76 (0.19)<br>WM:0.77 (0.11) |
| **BPF**<br>**(qMT)** | GM: 0.11 (0.01)<br>WM:0.10 (0.01) | GM: 0.11 (0.01)<br>WM:0.10 (0.01) | NA | NA | GM: 0.11 (0.01)<br>WM:0.10 (0.01) | GM: 0.11 (0.01)<br>WM:0.10 (0.01) |
| **k [1/s]**<br>**(qMT)** | GM: 1.73 (0.14)<br>WM:1.52 (0.07) | GM: 1.75 (0.13)<br>WM:1.53 (0.07) | NA | NA | GM: 1.75 (0.14)<br>WM:1.53 (0.08) | GM: 1.75 (0.13)<br>WM:1.54 (0.08) |
| **T1 [ms]**<br>**(IR)** | GM: 1108 (22)<br>WM:1131 (5) | GM: 1111 (22)<br>WM:1131 (5) | NA | GM: 1110 (22)<br>WM:1130 (5) | NA | GM: 1108 (21)<br>WM:1130 (6) |
| **T2 [ms]**<br>**(mTE)** | GM: 83.8 (5.9)<br>WM: 94.8 (2.9) | GM: 82.7 (5.6)<br>WM: 93.4 (3.1) | GM: 83.3 (5.8)<br>WM: 94.2 (2.9) | NA | NA | GM: 83.0 (6.2)<br>WM: 93.8 (2.8) |

**Table 6.** Median values in grey and white matter of qMRI metrics obtained with scanners from vendor 2 following different denoising strategies. Median values from different subjects and scans are pooled so that figures in the table report mean and standard deviation (in brackets) of across subjects and scans. In all cases, values of metrics obtained after denoising are less than 5% different from the values obtained with no denoising.

|  | No denoising | Individual denoising | Joint denoising<br>DWI-mTE |
| --- | --- | --- | --- |
| **FA (DWI)** | GM: 0.68 (0.02)<br>WM: 0.78 (0.01) | GM: 0.68 (0.02)<br>WM: 0.78 (0.01) | GM: 0.68 (0.02)<br>WM: 0.78 (0.01) |
| **MD [$\mu\text{m}^2/\text{ms}$] (DWI)** | GM: 0.99 (0.09)<br>WM: 1.07 (0.08) | GM: 0.98 (0.09)<br>WM: 1.07 (0.08) | GM: 0.98 (0.09)<br>WM: 1.07 (0.08) |
| **MK (DWI)** | GM: 0.83 (0.05)<br>WM: 0.83 (0.06) | GM: 0.81 (0.07)<br>WM: 0.83 (0.06) | GM: 0.82 (0.06)<br>WM: 0.83 (0.07) |
| **T2 [ms] (mTE)** | GM: 89.1 (2.7)<br>WM: 104.6 (1.2) | GM: 87.8 (2.7)<br>WM: 102.7 (2.0) | GM: 88.4 (2.7)<br>WM: 104.1 (1.9) |

Tables 7 and 8 report within-grey and within-white matter CoV for the various qMRI metrics and for different denoising strategies. Table 7 reports figures from vendor 1, while Table 8 from vendor 2 (data from both systems from vendor 2 pooled together). The tables show that MP-PCA denoising leads to reductions of CoV for various metrics of 5% or more compared to the case with no denoising, as for example for FA, MK, BPF and T_1_ for vendor 1 and MK for vendor 2. Some increases of CoV are observed (for example for MD in white matter for vendor 2). For vendor 1, the strongest reductions in CoV are observed for joint multimodal MP-PCA denoising.

**Table 7.** Percentage CoV in grey and white matter for the various qMRI metrics obtained from vendor 1 (London, UK) with different denoising strategies. The table reports CoV = 100 *iqr/median*, where *iqr* and *median* are respectively the interquartile range and the median of a metric within grey/white matter. CoV from different subjects and scans are pooled so that figures report mean and standard deviation (in brackets) of CoV across subjects and scans. Reductions in mean CoV with more than 5% as compared to the case with no denoising are labelled in bold font and grey shadowing (no increase of mean CoV greater than 5% is observed).

|  | No denoising | Individual denoising | Joint denoising DWI-mTE | Joint denoising DWI-IR | Joint denoising DWI-qMT | Joint denoising of all |
| --- | --- | --- | --- | --- | --- | --- |
| **FA CoV [%] (DWI)** | GM: 23.9 (3.3)<br>WM: 24.3 (2.8) | GM: 23.7 (3.3)<br>WM:24.2 (3.0) | GM: 23.6 (3.6)<br>WM:23.8 (3.1) | GM: 24.4 (4.5)<br>WM:23.8 (3.1) | GM: 23.2 (4.7)<br>WM: 23.2 (2.7) | **GM: 22.6 (4.6)**<br>WM:23.3 (2.8) |
| **MD CoV [%] (DWI)** | GM: 31.2 (15.5)<br>WM: 56.7 (3.3) | GM: 30.7 (14.4)<br>WM:58.3 (4.1) | GM: 30.7 (13.5)<br>WM: 58.0 (4.2) | GM: 30.5 (13.9)<br>WM:57.8 (4.1) | GM: 29.6 (15.0)<br>WM:57.6 (2.7) | **GM: 29.1 (15.8)**<br>WM:57.6 (3.4) |
| **MK CoV [%] (DWI)** | GM: 70.6 (65.5)<br>WM:58.9 (44.0) | **GM: 64.3 (59.6)**<br>**WM:53.4 (39.7)** | GM: 69.2 (76.6)<br>**WM:51.9 (37.9)** | GM: 71.1 (81.6)<br>**WM:52.7 (39.3)** | **GM: 61.7 (73.2)**<br>**WM:50.7 (39.1)** | **GM: 59.3 (70.0)**<br>**WM:50.9 (39.2)** |
| **BPF CoV [%] (qMT)** | GM: 44.5 (7.4)<br>WM:73.4 (12.3) | GM: 42.5 (8.7)<br>WM:72.5 (14.6) | NA | NA | **GM: 42.3 (8.1)**<br>WM:71.5 (15.7) | **GM: 40.9 (7.1)**<br>WM:70.9 (14.9) |
| **k CoV [%] (qMT)** | GM: 38.0 (9.9)<br>WM:75.6 (16.6) | GM: 38.8 (11.0)<br>WM:74.1 (15.6) | NA | NA | GM: 37.9 (10.4)<br>WM:73.5 (15.8) | GM: 37.1 (10.0)<br>WM:73.2 (15.8) |
| **T1 CoV [%] (IR)** | GM: 11.7 (6.9)<br>WM:41.7 (39.8) | **GM:10.9 (5.4)**<br>WM:41.5 (40.1) | NA | **GM:10.9 (6.3)**<br>WM:41.7 (40.2) | NA | **GM: 10.2 (6.6)**<br>WM:39.9 (40.6) |
| **T2 CoV [%] (mTE)** | GM:27.8 (9.5)<br>WM:44.4 (13.3) | GM:28.2 (10.2)<br>WM:44.6 (13.4) | GM: 27.5 (10.5)<br>WM:44.7 (13.8) | NA | NA | GM: 26.5 (6.8)<br>WM:43.7 (13.1) |

**Table 8.** Percentage CoV in grey and white matter for the various qMRI metrics obtained from vendor 2 (New York, NY, USA and Montreal, Canada) with different denoising strategies. The table reports CoV = 100 *iqr/median*, where *iqr* and *median* are respectively the interquartile range and the median of a metric within grey/white matter. CoV from different subjects and scans are pooled so that figures report mean and standard deviation (in brackets) of CoV across subjects and scans. Improvements of mean CoV greater than 5% compared to the case with no denoising (i.e. lower CoV) are labelled in bold font and grey shadowing (no worsening of mean CoV greater than 5% is observed).

|  | No denoising | Individual denoising | Joint denoising DWI-mTE |
| --- | --- | --- | --- |
| **FA CoV [%] (DWI)** | GM: 32.6 (1.0)<br>WM: 23.2 (1.4) | GM: 33.1 (1.7)<br>WM: 22.8 (2.0) | GM: 33.1 (2.8)<br>WM: 22.9 (1.9) |
| **MD CoV [%] (DWI)** | GM: 29.7 (7.1)<br>WM: 39.7 (4.1) | GM: 30.1 (7.3)<br>WM: 41.0 (3.4) | GM: 28.9 (4.7)<br>WM: 40.6 (4.9) |
| **MK CoV [%] (DWI)** | GM: 46.9 (7.8)<br>WM: 52.0 (6.7) | **GM: 39.7 (9.2)**<br>**WM: 44.1 (4.3)** | **GM: 38.9 (8.1)**<br>**WM: 44.3 (5.8)** |
| **T2 CoV [%] (mTE)** | GM: 26.3 (1.9)<br>WM: 39.4 (8.1) | GM: 25.3 (2.5)<br>WM: 40.1 (8.1) | GM: 26.5 (0.9)<br>WM: 39.7 (7.9) |

## 5. Discussion

### 5.1 Summary and key findings

This work demonstrates the advantages of multimodal qMRI of the spinal cord in vivo with unified MRI signal readout. The unified readout enables matching resolution and distortions across different contrasts, thereby also facilitating joint analyses and computational modelling of multi-contrast signal. These include MP-PCA denoising, a recent noise mitigation technique tailored for *redundant* qMRI protocols, i.e. such that the number of measurements is much higher than the underlying number of signal components. The unified readout enables efficient MP-PCA denoising of modalities that would exhibit little redundancy if denoised individually.

Our key findings are that a unified readout enables reliable and detailed microstructural characterisation of the human cervical spinal cord in clinical setting, providing metrics of relaxometry and diffusion as well as myelin-sensitive indices with matched resolution and distortions. Moreover, MP-PCA appears as a valid tool to improve the intrinsic quality of unified readout acquisitions, as supported by both in vivo and in silico data. Finally, this approach is feasible on 3T MRI systems from two major vendors.

### 5.2 In silico study

We have designed and run computer simulations to test whether a unified readout offers opportunities for MP-PCA denoising of qMRI modalities that exhibit limited redundancy (i.e. a number of measurements comparable to the number of signal generators), for which effective MP-PCA denoising remains challenging. For this purpose, we synthesised spinal cord qMRI data for a rich protocol consisting of DW, qMT, IR and mTE imaging, using anatomical information from the SCT atlas. We generated signals using literature values for microstructural parameters in MRI signal forward models, ensuring within-tissue variability to avoid obvious signal redundancy.

Our simulations suggest that a unified readout has indeed the potential of supporting more efficient MP-PCA denoising for modalities limited in redundancy, as for example mono-exponential and multi-exponential (Does et al., 2019) relaxation mapping. Denoising these modalities jointly with more redundant modalities enables more efficient noise mitigation in the former, thus suggesting that some of the information carried by the signal generators may be at least in part shared among MRI contrasts. Interestingly, joint multimodal denoising did not affect those modalities that are already redundant, as for example DW imaging, thus providing another piece of evidence supporting the idea that different contrasts may share some of the signal sources.

### 5.3 In vivo study

In this paper we tested multi-parametric qMRI of the spinal cord with unified readout using 3T MRI scanners from two major vendors (Philips and Siemens). We obtained a number of qMRI metrics that are promising biomarkers in several spinal cord conditions, including diffusion parameters FA, MD and MK; qMT two-pool BPF and exchange rate *k*; relaxation time constants T_1_ and T_2_. We studied whether MP-PCA improves the quality of raw MRI signals as well as quantitative maps, by evaluating distributions of signal residuals as well as within-tissue variability of metrics via a CoV.

Our multi-vendor data demonstrate the feasibility of implementing reliable multi-parametric qMRI of the spinal cord with unified readout. A unified readout provides matched resolution and distortions across MRI contrasts, and ensures comparability of signals across a rich set of qMRI measurements. Moreover, it enables the development of unified analysis pipelines, spanning from motion correction, to data denoising and potentially model fitting, paving the way to joint modelling of multi-contrast signals (Kim et al., 2017). Importantly, it may be useful in techniques that combine information from diffusion with relaxation/myelin-sensitive indices, as for example g-ratio MRI (Campbell et al., 2018; Duval et al., 2017; Stikov et al., 2015)), where matched EPI distortions (Irfanoglu et al., 2015) are crucial (Campbell et al., 2018). Our unified readout provides images with the same distortions across all contrasts and without significant losses of resolution for scans that would be typically performed with a different readout compared to DW imaging (Duval et al., 2017; Ljungberg et al., 2017), i.e. myelin mapping and relaxometry.

In this paper we also exploited synergies across MRI contrasts to evaluate the performance of MP-PCA denoising in the spinal cord. To our knowledge, this is the first time that the practical utilities of MP-PCA has been characterised in detail in the human spinal cord in vivo. Moreover, this is the first time that MP-PCA has been tested across that many MRI contrasts in a unified acquisition. Our analyses demonstrate that MP-PCA effectively mitigates noise in all modalities and for both vendors. Importantly, distributions of residuals show that the efficiency of MP-PCA in enhancing the quality of modalities with limited redundancy (i.e. IR and mTE imaging) can be improved by denoising these modalities jointly with more redundant schemes. Such results mirror findings in simulations, and thus suggest that some of the information about MRI signal sources may be shared across contrasts.

Our joint multimodal denoising relies on the hypothesis of noise homoscedasticity across MRI contrasts. Supplementary material S2 and S3 shows that the estimated noise level on modalities other than DWI follows the same trends as those of estimates from DWI in both simulations (S2) and in vivo (S3). Nonetheless, the two supplementary documents also demonstrate that estimating the noise level is a very challenging task: the estimates of the noise level are highly variable per se. Moreover, Supplementary material S3 also reveals systematic differences between noise standard deviation estimates from DW imaging compared to other modalities, such as qMT. This is likely attributed to the stronger departures from the hypothesis of Gaussian noise underlying MP-PCA in DW imaging, due to lower SNR and stronger noise floor effects (Koay and Basser, 2006), and to the fact that qMT suffers from stronger physiological noise that may resemble thermal noise (qMT is not cardiac gated). Nonetheless, it should be remembered that noise levels estimated on modalities with limited redundancy (e.g. multi-TE imaging) are not reliable, as the limited number of measurements does not allow the MP distribution to emerge, spoiling the detection of noisy eigenvalues in MP-PCA (Veraart et al., 2016b). Importantly, such differences in terms of noise level estimates among modalities introduce little to no bias in downstream quantitative parameter maps, and therefore do not appear to be a concerning issue for practical MP-PCA deployment.

We also investigated the effect of MP-PCA denoising on the quality of popular parametric maps. To this end, we studied median values of metrics within grey/white matter as well as metric variabilities as quantified by a CoV. Our experiments show that MP-PCA introduces little to no biases in any of the metrics, irrespectively of the chosen denoising strategy (joint multimodal denoising vs modality-wise denoising). The difference in median values between metrics obtained with denoising compared to the case with no denoising are ±5 % or less. These differences, which are very low, likely reflect the intrinsic susceptibility of the different model fitting routines to noise fluctuations and noise floors, and are therefore expected since noise-floor mitigation (Koay and Basser, 2006) was performed following MP-PCA denoising. Conversely, MP-PCA does decrease metric variability, at it leads to tissue-wise CoV reduction much higher 5% for many metrics, as for example for MK (both vendors), FA, BPF and T_1_ (vendor 1). The reduction in variability is the highest for metrics like MK, which carry important information about tissue microstructure, and that are notoriously difficult to estimate (Veraart et al., 2011).

Our parametric maps follow known trends and contrasts, with some differences in terms of relaxometry metrics, e.g. low contrast white/grey matter contrast for T1 and T2. This difference may be explained by residual CSF pulsation that corrupts neighbouring white matter signals, and by the fact that literature values for T1 and T2 are typically obtained with different readout strategies compared to the employed single-shot EPI (Smith et al., 2008). Another explanation, especially for vendor 1, may be related to the coarse resolution of the anatomical scan, required to limit scan time, as this was used for grey matter segmentation potentially introducing partial volume effects in the tissue masks. Overall, while grey/white matter contrasts in parametric maps are similar in data from both vendors, systematic differences between metric values (Table 5 vs 6) and variability (Table 7 vs 8) are seen. Several factors may have contributed to these differences between vendors, namely in: intrinsic SNRs; rFOV techniques; resolution of the anatomical scan used for grey/white matter segmentation; parallel imaging/reconstruction techniques; qMRI protocol; between-subject biological differences.

Finally, we point out that we took care to use the same registration transformations to correct for motion in all denoising strategies, estimating motion on the non-denoised data. We followed this motion correction strategy on purpose, as our focus was to study the effects of thermal noise mitigation on parametric maps. It is possible that the benefits of MP-PCA may extend beyond thermal noise mitigation and may also improve post-processing such as motion correction, as shown in other studies (Ades-Aron et al., 2018), which will be the subject of future investigations.

### 5.4 Considerations and future directions

We acknowledge a number of limitations of our approach.

Firstly, our unified protocol was more comprehensive on the system from vendor 1 as compared to vendor 2. This was due to the practical availability of MRI sequences at the time of acquisition of the data. In future we plan to expand our protocol on vendor 2 to better characterise MP-PCA performance across contrasts.

Secondly, the DW imaging protocol for the vendor 2 differed between the system in New York and the one in Montreal, with the latter being slightly longer. This was due to a choice in the design of the protocol in Montreal, which would enable the inclusion of the scan in other ongoing group studies. Nonetheless, the acquisition suffices to demonstrate the potential of a unified readout and to explore multimodal denoising.

Furthermore, in the future we plan to expand our sample size to better characterise the potential of our unified acquisition and denoising in real clinical settings on patients as, for example, in multiple sclerosis.

### 5.5 Conclusions

Multi-parametric qMRI of the spinal cord with unified readout (i.e. with matched resolution and distortions) is advantageous and provides robust microstructural metrics sensitive to axonal characteristics, relaxometry and myelin. Our unified acquisition paves the way to joint modelling of multi-contrast signals, and offers unique opportunities for image quality enhancement with techniques such as MP-PCA denoising, proven here to be a useful pre-processing step in spinal cord MRI analysis pipelines.

## Acknowledgements

The authors thank all the volunteers for their time, Mary Bruno and Alexandru Foias for their support with data acquisition, and Manuel J. Cardoso for mentoring.

## Author contribution statement

FG: Conceptualization; Methodology; Software; Formal analysis; Investigation; Data Curation; Writing -Original Draft; Visualisation; Funding acquisition; Project administration. MB, JV, TS: Conceptualization; Methodology; Software; Investigation; Writing - Review & Editing. JCA, TMS, DCA, DSN, EF, CGWK: Conceptualization; Methodology; Resources, Supervision, Funding acquisition; Writing - Review & Editing.

## Data and code availability statement

The synthetic spinal cord phantom is made freely available online (http://github.com/fragrussu/PaperScripts/tree/master/sc_unireadout/sc_phantom). All scripts written to analyse the data are also made openly available (http://github.com/fragrussu/PaperScripts/tree/master/sc_unireadout). The Python routines for relaxometry and DKI fitting are made freely available online as part of the open-access GitHub repositories MyRelax (http://github.com/fragrussu/MyRelax) and MRItools (http://github.com/fragrussu/MRItools). All relevant third party dependencies are clearly indicated.

The in vivo data cannot be made openly available online due to privacy issues of clinical data according to GDPR regulations. Researchers interested in accessing the in vivo data from vendor 1 can contact Prof Claudia Gandini Wheeler-Kingshott. A data sharing agreement enabling academic and research use will be stipulated. Researchers interested in accessing the in vivo data from vendor 2 can contact: Prof Timothy Shepherd for the New York data; Prof Julien Cohen-Adad for the Montreal data. A data sharing agreement will be stipulated with either New York University, Polytechnique Montreal or both, enabling academic and research use.

Other third party code used in this project is freely available online. This includes: MP-PCA denoising, SCT, NiftyReg, NiftK, DiPy and FSL (MP-PCA: http://github.com/NYU-DiffusionMRI/mppca_denoise; SCT: http://github.com/neuropoly/spinalcordtoolbox; NiftyReg: http://cmictig.cs.ucl.ac.uk/wiki/index.php/NiftyReg; NiftyK: http://github.com/NifTK/NifTK; DiPy: http://dipy.org; FSL: http://fsl.fmrib.ox.ac.uk/fsl/fslwiki).

The code for qMT signal synthesis and analysis (two-pool fitting) is not openly available online. Researchers interested in accessing the code can contact Dr Marco Battiston, who would release a copy under a license/sharing agreement enabling academic and research use.

## Supplementary material S1: the two-pool MT model

The two-pool magnetisation transfer (MT) model is used in this paper in both simulations and in vivo analyses. The model describes the evolution of the magnetisation of two exchanging ^1^H pools, i.e. free protons in bulk water and protons bound to macromolecules (Henkelman et al., 1993), in presence of off-resonance irradiation *b*_1,off_(*t*) with carrier frequency *f*_0_ + Δ*f*_*c*_ (with Δ*f*_*c*_ being the offset with respect to water resonance frequency *f*_0_), assumed to be played out at times 0 ≤ *t* ≤ *t*_off_. On-resonance excitation is then assumed to be played out at *t* > *t*_off_, after a generic delay *t*_delay_.

Let us indicate with 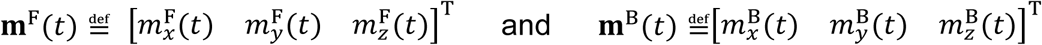 the free and bound proton magnetisations during the off-resonance irradiation, and with 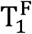 and 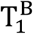 and with 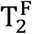 and 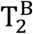 their respective longitudinal and transverse relaxation times (Portnoy and Stanisz, 2007). Let us also indicate with *k* the exchange rate between free to bound protons, with BPF the bound pool fraction (fraction of protons belonging to the bound pool) and with *γ* the proton gyromagnetic ratio 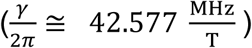.

The MT-weighting factor ***w*** in Eq. 5 (main manuscript) is defined as the value of the free pool longitudinal magnetisation at the end of off-resonance irradiation, i.e.:

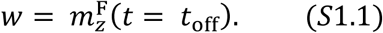

The value of *w* in Eq. S1.1 is obtained directly by numerical integration of the two-pool Bloch equations, which are written as:

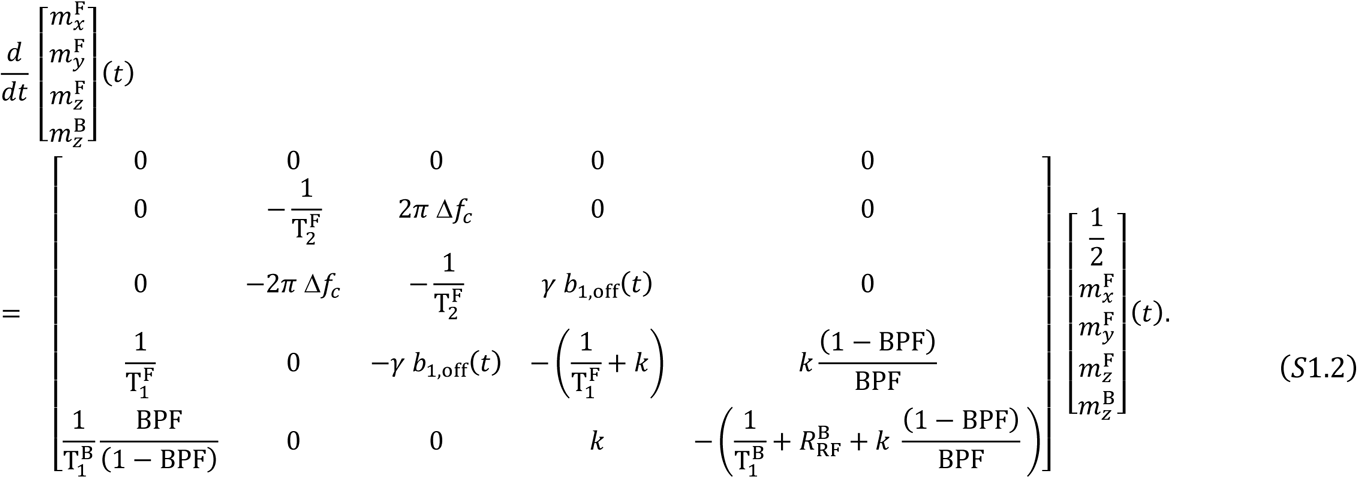

For the integration of equation S1.2, it is assumed that 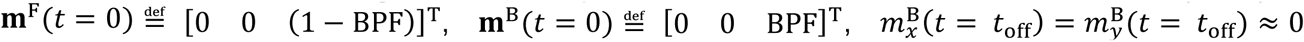 and that the bound pool off-resonance absorption can be modelled based on a super-Lorentzian line shape. Under this assumption, the absorption term 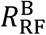 is a function of 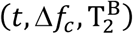 and is modelled as (Portnoy and Stanisz, 2007):

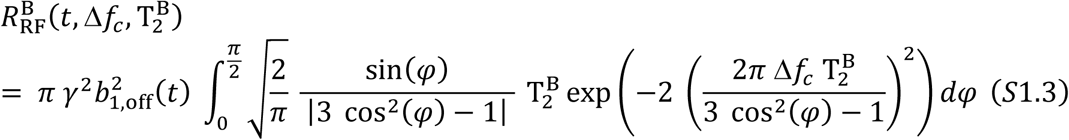

For the practical integration of Eq. S1.2, it is assumed that 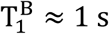 (Battiston et al., 2018a), and that the free pool longitudinal relaxation time (i.e. 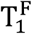) is linked to the observable T_1_ relaxation time as (Portnoy and Stanisz, 2007):

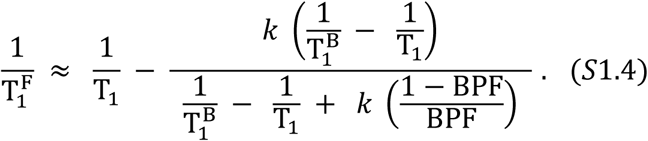

## Supplementary material S2: estimated noise levels in simulations

In this supplementary material we compare the estimates of noise levels provided by MP-PCA denoising when denoising different modalities in simulations.

In the following supplementary figures, we scatter the estimated noise standard deviation obtained from denoising the DWI against the values obtained by denoising the other modalities, either alone or jointly with DWI.

**Fig. S2.1.**
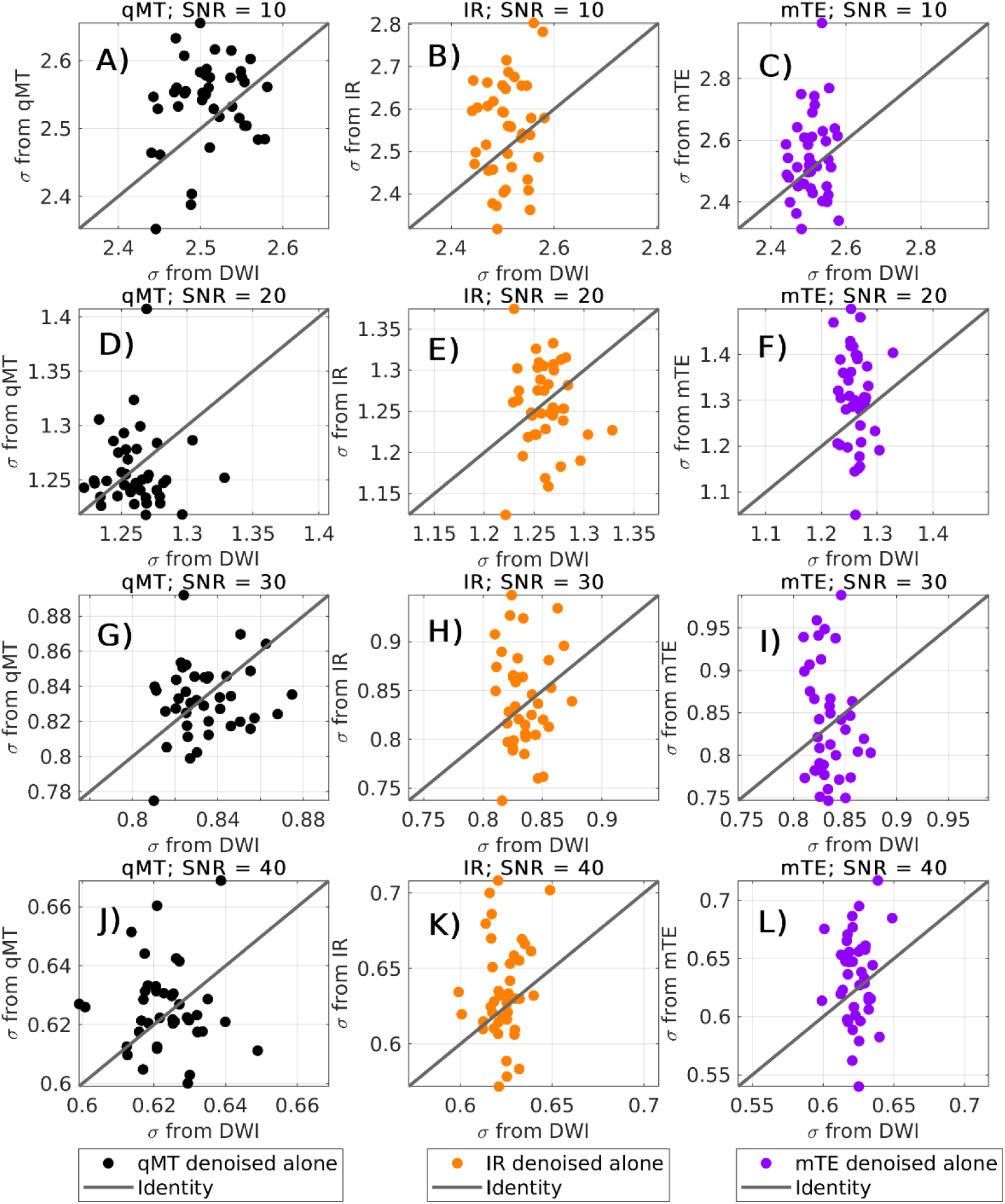
Estimated noise standard deviation *σ* obtained denoising each modality independently. Modalities are: diffusion-weighted imaging (DWI); quantitative magnetisation transfer (qMT); inversion recovery (IR); multi-echo time (mTE). Each plot scatters values of *σ* obtained from qMT (left), IR (middle) or mTE (right), reported on the y-axis, against *σ* provided by denoising DWI alone (x-axis). Different rows shows different signal-to-noise ratio (SNR) levels (from top to bottom: SNR of 10, 20, 30 and 40).

**Fig. S2.2.**
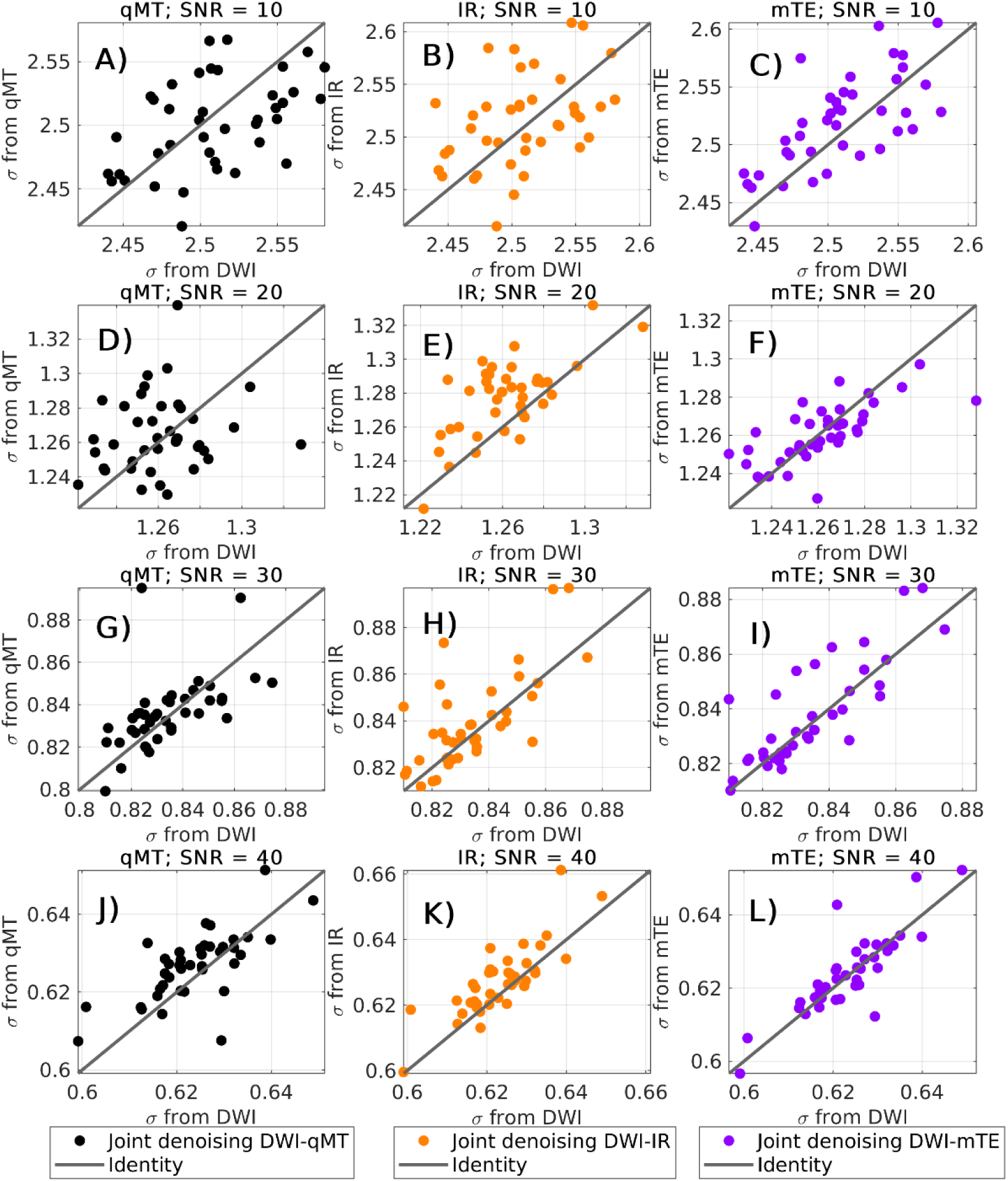
Estimated noise standard deviation *σ* obtained denoising each of quantitative magnetisation transfer (qMT), inversion recovery (IR) and multi-echo time (mTE) jointly with diffusion-weighted imaging (DWI). Each plot scatters values of *σ* obtained from joint denoising DWI-qMT (left), DWI-IR (middle) or DWI-mTE (right), reported on the y-axis, against *σ* provided by denoising DWI alone (x-axis). Different rows shows different signal-to-noise ratio (SNR) levels (from top to bottom: SNR of 10, 20, 30 and 40).

## Supplementary material S3: estimated noise levels in vivo

In this supplementary material we compare the estimates of noise levels provided by MP-PCA denoising on different modalities (in vivo data).

In the following supplementary figures, we scatter the estimated noise standard deviation obtained from denoising the DWI against the values obtained by denoising the other modalities, either alone or jointly with DWI. Moreover, we report the scatter plots with and without noise floor mitigation, and for the two vendors.

Results are shown in both cases when denoised images and estimated noise level are corrected for Rician noise bias with the method of moments [1] and whey they are not.

**Fig. S3.1.**
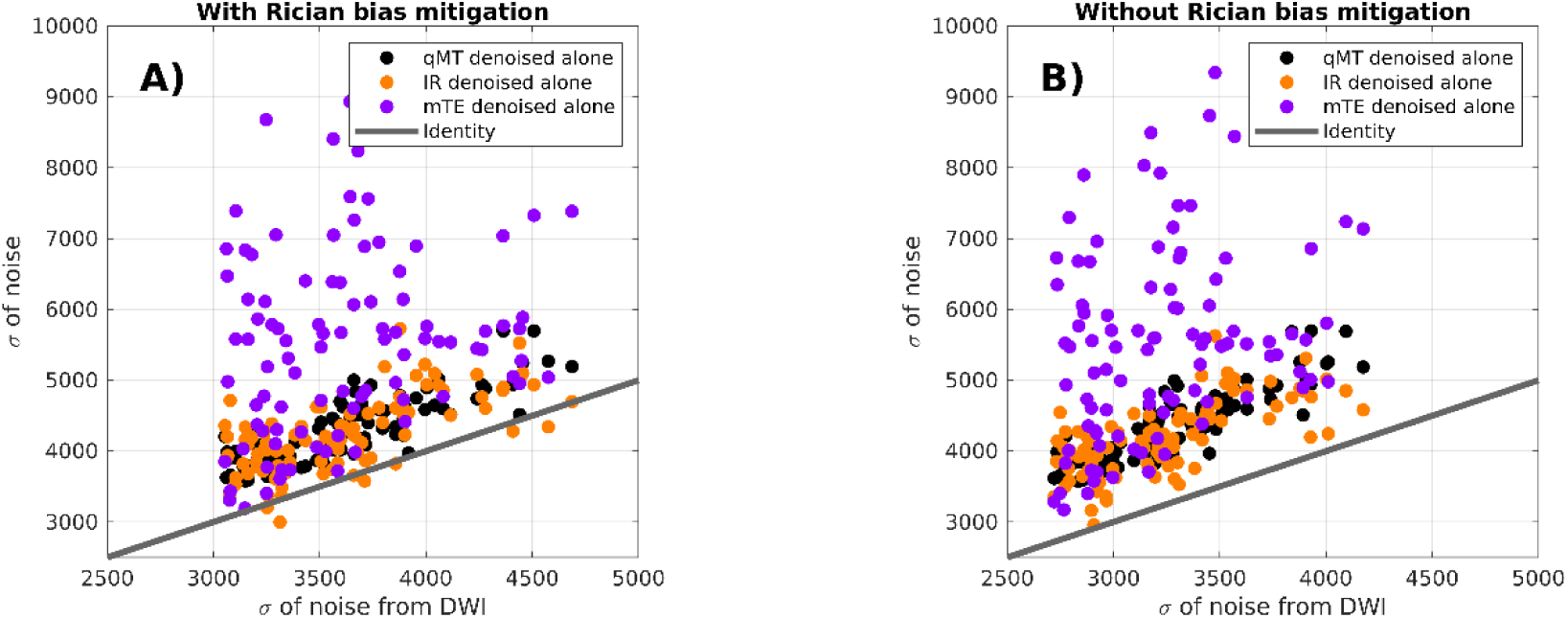
Estimated noise standard deviation for vendor 1 (Philips Achieva) obtained denoising each modality (qMT, IR and mTE) independently. Values are scattered against the noise level estimated from the DWI alone. In panel A) (left), the estimated noise level has been corrected for Rician bias, while in panel B) (to the right) this correction has not been performed. When generating the plots, data from all scans of all subjects have been pooled together.

**Figure S3.2.**
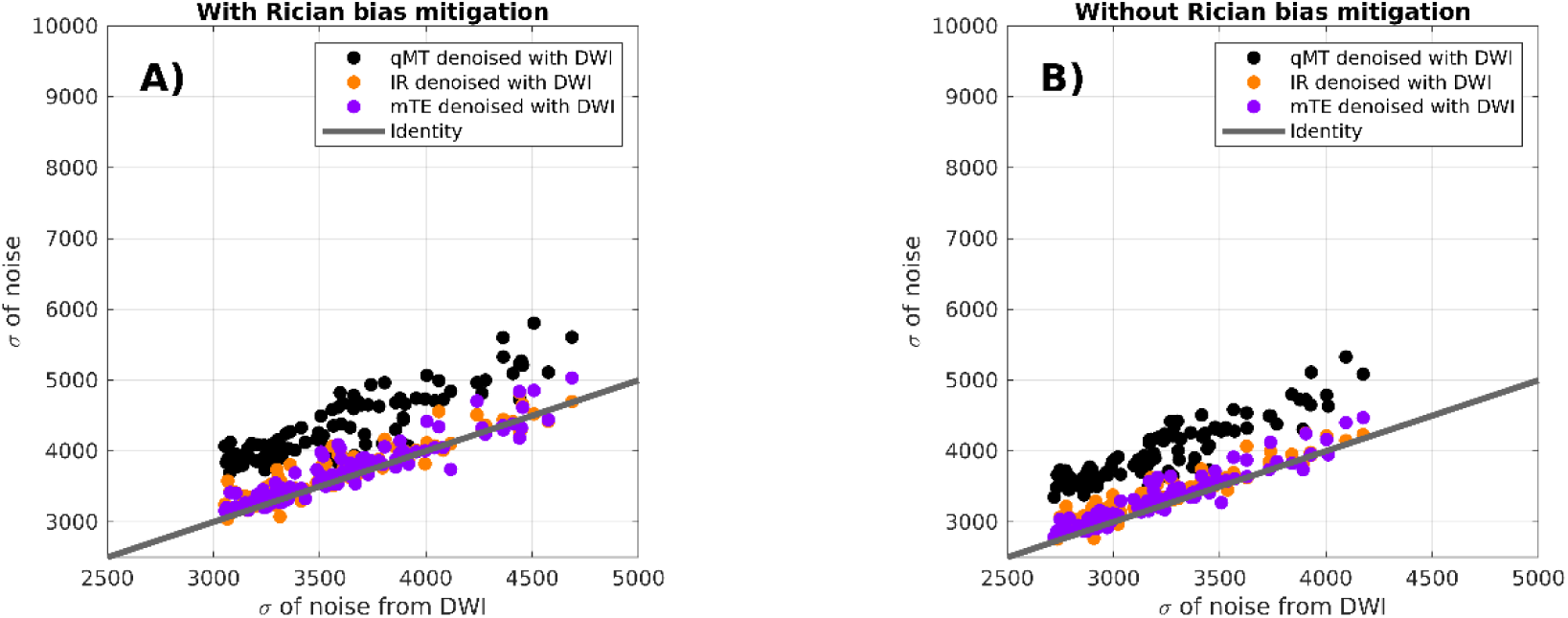
Estimated noise standard deviation for vendor 1 (Philips Achieva) obtained denoising each modality (qMT, IR and mTE) jointly with DWI. Values are scattered against the noise level estimated from the DWI alone. In panel A) (left), the estimated noise level has been corrected for Rician bias, while in panel B) (to the right) this correction has not been performed. When generating the plots, data from all scans of all subjects have been pooled together.

**Fig. S3.3.**
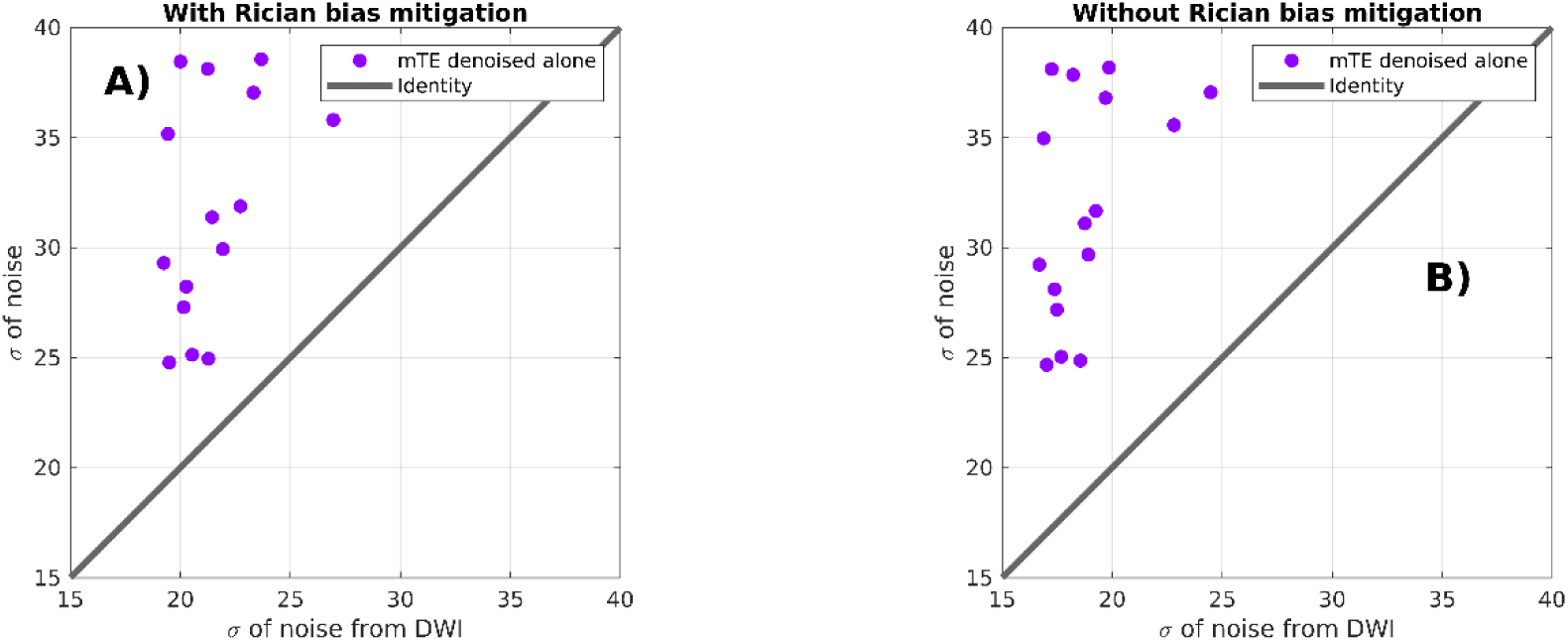
Estimated noise standard deviation for vendor 2 (two Siemens Prisma located in New York, USA and Montreal, Canada) obtained denoising mTE independently from DWI. Values are scattered against the noise level estimated from the DWI alone. In panel A) (left), the estimated noise level has been corrected for Rician bias, while in panel B) (to the right) this correction has not been performed. When generating the plots, data from all scans of all systems have been pooled together.

**Fig. S3.4.**
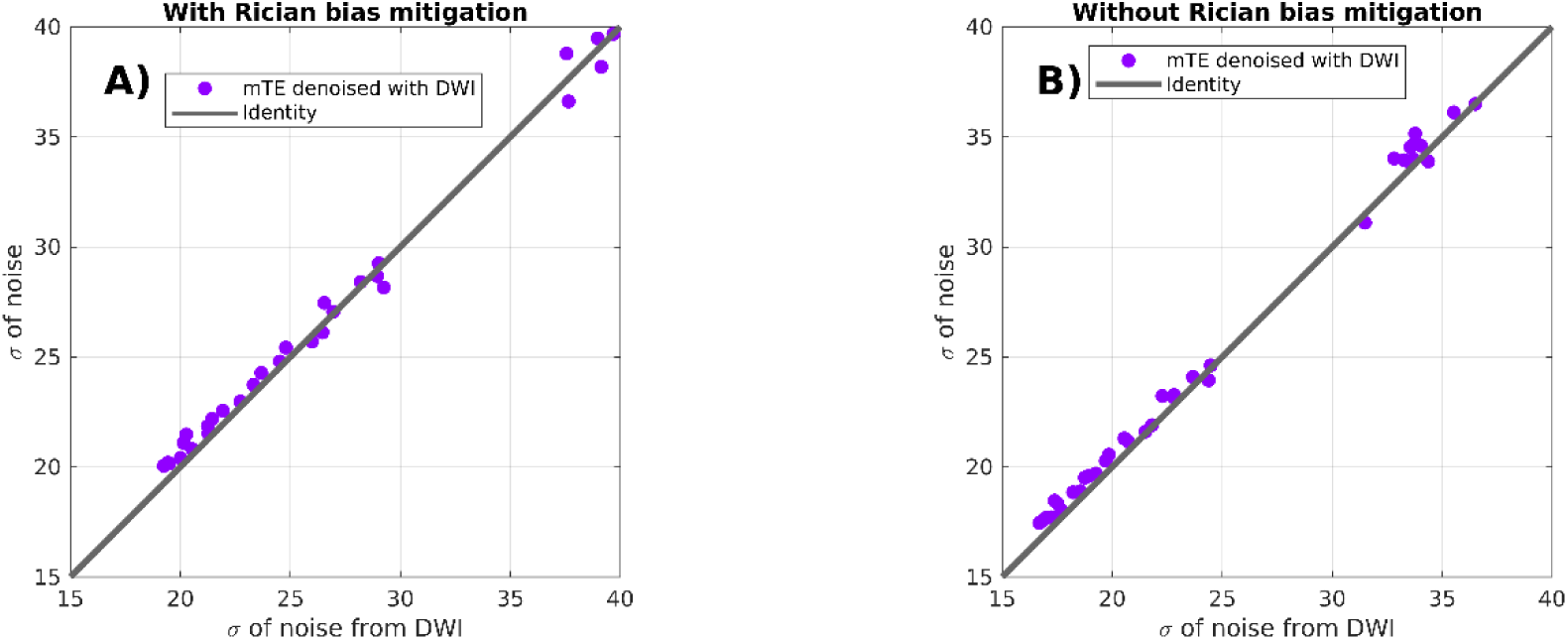
Estimated noise standard deviation for vendor 2 (two Siemens Prisma located in New York, USA and Montreal, Canada) obtained denoising mTE jointly with DWI. Values are scattered against the noise level estimated from the DWI alone. In panel A) (left), the estimated noise level has been corrected for Rician bias, while in panel B) (to the right) this correction has not been performed. When generating the plots, data from all scans of all systems have been pooled together.

## References

Ades-Aron, B., Veraart, J., Kochunov, P., McGuire, S., Sherman, P., Kellner, E., Novikov, D.S., Fieremans, E., 2018. Evaluation of the accuracy and precision of the diffusion parameter EStImation with Gibbs and NoisE removal pipeline. NeuroImage 183, 532–543.

Ahuja, C.S., Wilson, J.R., Nori, S., Kotter, M.R., Druschel, C., Curt, A., Fehlings, M.G., 2017. Traumatic spinal cord injury. Nature reviews Disease primers 3, 17018.

Barry, R.L., Vannesjo, S.J., By, S., Gore, J.C., Smith, S.A., 2018. Spinal cord MRI at 7T. NeuroImage 168, 437–451.

Basser, P.J., Mattiello, J., LeBihan, D., 1994. MR diffusion tensor spectroscopy and imaging. Biophysical journal 66, 259–267.

Battiston, M., Grussu, F., Ianus, A., Schneider, T., Prados, F., Fairney, J., Ourselin, S., Alexander, D.C., Cercignani, M., Gandini Wheeler-Kingshott, C.A., 2018a. An optimized framework for quantitative magnetization transfer imaging of the cervical spinal cord in vivo. Magnetic resonance in medicine 79, 2576–2588.

Battiston, M., Schneider, T., Prados, F., Grussu, F., Yiannakas, M.C., Ourselin, S., Gandini Wheeler-Kingshott, C.A., Samson, R.S., 2018b. Fast and reproducible in vivo T1 mapping of the human cervical spinal cord. Magnetic resonance in medicine 79, 2142–2148.

By, S., Xu, J., Box, B.A., Bagnato, F.R., Smith, S.A., 2017. Application and evaluation of NODDI in the cervical spinal cord of multiple sclerosis patients. NeuroImage: Clinical 15, 333–342.

By, S., Xu, J., Box, B.A., Bagnato, F.R., Smith, S.A., 2018. Multi-compartmental diffusion characterization of the human cervical spinal cord in vivo using the spherical mean technique. NMR in Biomedicine 31, e3894.

Campbell, J.S., Leppert, I.R., Narayanan, S., Boudreau, M., Duval, T., Cohen-Adad, J., Pike, G.B., Stikov, N., 2018. Promise and pitfalls of g-ratio estimation with MRI. NeuroImage 182, 80–96.

Cercignani, M., Bouyagoub, S., 2018. Brain microstructure by multi-modal MRI: Is the whole greater than the sum of its parts? NeuroImage 182, 117–127.

Ciccarelli, O., Cohen, J.A., Reingold, S.C., Weinshenker, B.G., Amato, M.P., Banwell, B., Barkhof, F., Bebo, B., Becher, B., Bethoux, F., 2019. Spinal cord involvement in multiple sclerosis and neuromyelitis optica spectrum disorders. The Lancet Neurology 18, 185–197.

Cohen-Adad, J., 2018. Microstructural imaging in the spinal cord and validation strategies. NeuroImage 182, 169–183.

De Leener, B., Kadoury, S., Cohen-Adad, J., 2014. Robust, accurate and fast automatic segmentation of the spinal cord. NeuroImage 98, 528–536.

De Leener, B., Lévy, S., Dupont, S.M., Fonov, V.S., Stikov, N., Collins, D.L., Callot, V., Cohen-Adad, J., 2017. SCT: Spinal Cord Toolbox, an open-source software for processing spinal cord MRI data. NeuroImage 145, 24–43.

De Santis, S., Barazany, D., Jones, D.K., Assaf, Y., 2016. Resolving relaxometry and diffusion properties within the same voxel in the presence of crossing fibres by combining inversion recovery and diffusion-weighted acquisitions. Magnetic resonance in medicine 75, 372–380.

Does, M.D., Olesen, J.L., Harkins, K.D., Serradas-Duarte, T., Gochberg, D.F., Jespersen, S.N., Shemesh, N., 2019. Evaluation of principal component analysis image denoising on multi-exponential MRI relaxometry. Magnetic resonance in medicine 81, 3503–3514.

Duval, T., Lévy, S., Stikov, N., Campbell, J., Mezer, A., Witzel, T., Keil, B., Smith, V., Wald, L.L., Klawiter, E., 2017. g-Ratio weighted imaging of the human spinal cord in vivo. NeuroImage 145, 11–23.

Duval, T., McNab, J.A., Setsompop, K., Witzel, T., Schneider, T., Huang, S.Y., Keil, B., Klawiter, E.C., Wald, L.L., Cohen-Adad, J., 2015. In vivo mapping of human spinal cord microstructure at 300 mT/m. NeuroImage 118, 494–507.

Grussu, F., Ianuş, A., Tur, C., Prados, F., Schneider, T., Kaden, E., Ourselin, S., Drobnjak, I., Zhang, H., Alexander, D.C., 2019. Relevance of time-dependence for clinically viable diffusion imaging of the spinal cord. Magnetic resonance in medicine 81, 1247–1264.

Grussu, F., Schneider, T., Zhang, H., Alexander, D.C., Wheeler–Kingshott, C.A., 2015. Neurite orientation dispersion and density imaging of the healthy cervical spinal cord in vivo. NeuroImage 111, 590–601.

Gudbjartsson, H., Patz, S., 1995. The Rician distribution of noisy MRI data. Magnetic resonance in medicine 34, 910–914.

Hendrix, P., Griessenauer, C.J., Cohen-Adad, J., Rajasekaran, S., Cauley, K.A., Shoja, M.M., Pezeshk, P., Tubbs, R.S., 2015. Spinal diffusion tensor imaging: a comprehensive review with emphasis on spinal cord anatomy and clinical applications. Clinical Anatomy 28, 88–95.

Henkelman, R.M., Huang, X., Xiang, Q.S., Stanisz, G., Swanson, S.D., Bronskill, M.J., 1993. Quantitative interpretation of magnetization transfer. Magnetic resonance in medicine 29, 759–766.

Irfanoglu, M.O., Modi, P., Nayak, A., Hutchinson, E.B., Sarlls, J., Pierpaoli, C., 2015. DR-BUDDI (Diffeomorphic Registration for Blip-Up blip-Down Diffusion Imaging) method for correcting echo planar imaging distortions. NeuroImage 106, 284–299.

Jensen, J.H., Helpern, J.A., Ramani, A., Lu, H., Kaczynski, K., 2005. Diffusional kurtosis imaging: the quantification of non-gaussian water diffusion by means of magnetic resonance imaging. Magnetic resonance in medicine 53, 1432–1440.

Jezzard, P., Balaban, R.S., 1995. Correction for geometric distortion in echo planar images from B0 field variations. Magnetic resonance in medicine 34, 65–73.

Johnstone, I.M., 2006. High dimensional statistical inference and random matrices. arXiv preprint math, 0611589.

Kearney, H., Miller, D.H., Ciccarelli, O., 2015. Spinal cord MRI in multiple sclerosis— diagnostic, prognostic and clinical value. Nature Reviews Neurology 11, 327.

Kim, D., Doyle, E.K., Wisnowski, J.L., Kim, J.H., Haldar, J.P., 2017. Diffusion-relaxation correlation spectroscopic imaging: a multidimensional approach for probing microstructure. Magnetic resonance in medicine 78, 2236–2249.

Koay, C.G., Basser, P.J., 2006. Analytically exact correction scheme for signal extraction from noisy magnitude MR signals. Journal of magnetic resonance 179, 317–322.

Lemberskiy, G., Fieremans, E., Veraart, J., Rosenkrantz, A., Novikov, D.S., 2018. Characterization of prostate microstructure using water diffusion and NMR relaxation. Frontiers in physics 6, 91.

Lévy, S., Benhamou, M., Naaman, C., Rainville, P., Callot, V., Cohen-Adad, J., 2015. White matter atlas of the human spinal cord with estimation of partial volume effect. NeuroImage 119, 262–271.

Ljungberg, E., Vavasour, I., Tam, R., Yoo, Y., Rauscher, A., Li, D.K., Traboulsee, A., MacKay, A., Kolind, S., 2017. Rapid myelin water imaging in human cervical spinal cord. Magnetic resonance in medicine 78, 1482–1487.

Lorenzi, R.M., Palesi, F., Castellazzi, G., Vitali, P., Anzalone, N., Bernini, S., Sinforiani, E., Micieli, G., Costa, A., D’Angelo, E., 2019. Unsuspected involvement of spinal cord in Alzheimer Disease. bioRxiv 10.1101, 673350v673353.

Massire, A., Taso, M., Besson, P., Guye, M., Ranjeva, J.-P., Callot, V., 2016. High-resolution multi-parametric quantitative magnetic resonance imaging of the human cervical spinal cord at 7T. NeuroImage 143, 58–69.

Mezer, A., Yeatman, J.D., Stikov, N., Kay, K.N., Cho, N.-J., Dougherty, R.F., Perry, M.L., Parvizi, J., Hua, L.H., Butts-Pauly, K., 2013. Quantifying the local tissue volume and composition in individual brains with magnetic resonance imaging. Nature medicine 19, 1667.

Modat, M., Ridgway, G.R., Taylor, Z.A., Lehmann, M., Barnes, J., Hawkes, D.J., Fox, N.C., Ourselin, S., 2010. Fast free-form deformation using graphics processing units. Computer methods and programs in biomedicine 98, 278–284.

Morozov, D., Rios, N.L., Duval, T., Foias, A., Cohen-Adad, J., 2018. Effect of cardiac-related translational motion in diffusion MRI of the spinal cord. Magnetic resonance imaging 50, 119–124.

Ning, L., Gagoski, B., Szczepankiewicz, F., Westin, C.-F., Rathi, Y., 2019. Joint RElaxation-Diffusion Imaging Moments (REDIM) to probe neurite microstructure. bioRxiv 10.1101, 598375v598371.

Novikov, D.S., Kiselev, V.G., Jespersen, S.N., 2018. On modeling. Magnetic resonance in medicine 79, 3172–3193.

Portnoy, S., Stanisz, G.J., 2007. Modeling pulsed magnetization transfer. Magnetic resonance in medicine 58, 144–155.

Rieseberg, S., Frahm, J., Finsterbusch, J., 2002. Two-dimensional spatially-selective RF excitation pulses in echo-planar imaging. Magnetic resonance in medicine 47, 1186–1193.

Schilling, K.G., By, S., Feiler, H.R., Box, B.A., O’Grady, K.P., Witt, A., Landman, B.A., Smith, S.A., 2019. Diffusion MRI microstructural models in the cervical spinal cord–Application, normative values, and correlations with histological analysis. NeuroImage 201, 116026.

Slator, P.J., Hutter, J., Palombo, M., Jackson, L.H., Ho, A., Panagiotaki, E., Chappell, L.C., Rutherford, M.A., Hajnal, J.V., Alexander, D.C., 2019. Combined diffusion-relaxometry MRI to identify dysfunction in the human placenta. Magnetic resonance in medicine 82, 95–106.

Smith, S.A., Edden, R.A., Farrell, J.A., Barker, P.B., Van Zijl, P.C., 2008. Measurement of T1 and T2 in the cervical spinal cord at 3 tesla. Magnetic resonance in medicine 60, 213–219.

Stikov, N., Campbell, J.S., Stroh, T., Lavelée, M., Frey, S., Novek, J., Nuara, S., Ho, M.-K., Bedell, B.J., Dougherty, R.F., 2015. In vivo histology of the myelin g-ratio with magnetic resonance imaging. NeuroImage 118, 397–405.

Stroman, P.W., Wheeler-Kingshott, C., Bacon, M., Schwab, J., Bosma, R., Brooks, J., Cadotte, D., Carlstedt, T., Ciccarelli, O., Cohen-Adad, J., 2014. The current state-of-the-art of spinal cord imaging: methods. NeuroImage 84, 1070–1081.

Summers, P., Stämpfli, P., Jaermann, T., Kwiecinski, S., Kollias, S., 2006. A preliminary study of the effects of trigger timing on diffusion tensor imaging of the human spinal cord. American Journal of Neuroradiology 27, 1952–1961.

Taso, M., Girard, O.M., Duhamel, G., Le Troter, A., Feiweier, T., Guye, M., Ranjeva, J.P., Callot, V., 2016. Tract-specific and age-related variations of the spinal cord microstructure: a multi-parametric MRI study using diffusion tensor imaging (DTI) and inhomogeneous magnetization transfer (ihMT). NMR in Biomedicine 29, 817–832.

van Es, M.A., Hardiman, O., Chio, A., Al-Chalabi, A., Pasterkamp, R.J., Veldink, J.H., Van den Berg, L.H., 2017. Amyotrophic lateral sclerosis. The Lancet 390, 2084–2098.

Vannesjo, S.J., Miller, K.L., Clare, S., Tracey, I., 2018. Spatiotemporal characterization of breathing-induced B0 field fluctuations in the cervical spinal cord at 7T. NeuroImage 167, 191–202.

Veraart, J., Fieremans, E., Novikov, D.S., 2016a. Diffusion MRI noise mapping using random matrix theory. Magnetic resonance in medicine 76, 1582–1593.

Veraart, J., Novikov, D.S., Christiaens, D., Ades-Aron, B., Sijbers, J., Fieremans, E., 2016b. Denoising of diffusion MRI using random matrix theory. NeuroImage 142, 394–406.

Veraart, J., Novikov, D.S., Fieremans, E., 2018. TE dependent Diffusion Imaging (TEdDI) distinguishes between compartmental T2 relaxation times. NeuroImage 182, 360–369.

Veraart, J., Van Hecke, W., Sijbers, J., 2011. Constrained maximum likelihood estimation of the diffusion kurtosis tensor using a Rician noise model. Magnetic resonance in medicine 66, 678–686.

Verma, T., Cohen-Adad, J., 2014. Effect of respiration on the B0 field in the human spinal cord at 3T. Magnetic resonance in medicine 72, 1629–1636.

Wheeler-Kingshott, C., Stroman, P.W., Schwab, J., Bacon, M., Bosma, R., Brooks, J., Cadotte, D., Carlstedt, T., Ciccarelli, O., Cohen-Adad, J., 2014. The current state-of-the-art of spinal cord imaging: applications. NeuroImage 84, 1082–1093.

Wheeler-Kingshott, C.A., Parker, G.J., Symms, M.R., Hickman, S.J., Tofts, P.S., Miller, D.H., Barker, G.J., 2002. ADC mapping of the human optic nerve: increased resolution, coverage, and reliability with CSF-suppressed ZOOM-EPI. Magnetic resonance in medicine 47, 24–31.

Yarnykh, V.L., 2007. Actual flip-angle imaging in the pulsed steady state: a method for rapid three-dimensional mapping of the transmitted radiofrequency field. Magnetic resonance in medicine 57, 192–200.

